# Three pollen QTLs are sufficient to partially break stylar unilateral incompatibility of *Solanum pennellii*

**DOI:** 10.1101/2024.04.23.590840

**Authors:** Wenchun Ma, Yumei Li, Mingxia He, Ian T. Baldwin, Han Guo

## Abstract

● In tomato clade, both <u>s</u>elf-incompatibility(*S*-) locus-dependent and –independent mechanisms are present in unilateral incompatibility (UI). Several stylar UI factors mediating *S*-locus-independent mechanism have been gradually uncovered, however, related pollen UI (*pui*) factors still await more studies.
● Quantitative trait loci (QTL) mapping, comparative transcriptomics and introgression lines (ILs) /inbred backcross lines (BILs)-based functional analysis were applied for identification and functional validation of *pui* QTLs between self-compatible (SC) *Solanum lycopersicum* and SC *S. pennellii* LA0716.
● In addition to the reported *pui10.1* (*SpFPS2*), two *pui* QTLs (*pui6.2* and *pui12.1*) were identified. Pollen tubes of cultivated tomatoes introgressed with three *pui* loci could partially break the stylar UI response in LA0716 styles. Furthermore, a conserved function of *pui12.1* was found in the styles of SC *S*. *habrochaites* LA0407 and SC *S*. *chmielewskii* LA1028. Three candidate genes linked to *pui6.2* and *pui12.1* were screened for further analysis.
● This study reveals a potential synergistic relationship of the three *pui* QTLs in breaking stylar UI response of LA0716 and will advance our understanding of the genetic mechanisms underlying UI in the tomato clade.

**Highlight:** Our data revealed that breaking the stylar unilateral incompatibility of *Solanum pennellii* LA0716 requires not only the reported pollen UI (*pui*) factor but also another two *pui* QTLs identified in this study.

## Introduction

In flowering plants, multiple reproductive barriers (RBs) have evolved that restrict gene flow (Rieseberg & Willis, 2007, Lowry *et al*., 2008) with both ecological (e.g. maintenance of genetic diversity in populations by intraspecific self-incompatibility (Goldberg *et al*., 2010)) and evolutionary (e.g. reproductive isolation by interspecific RBs (Moyle *et al*., 2004, Bedinger *et al*., 2011)) consequences. In the interactions between pollen and pistils, various post-pollination and pre-zygotic barriers have evolved (Hiscock & Allen, 2008, Broz & Bedinger, 2021), which can mediate either intraspecific (e.g. self-incompatibility, SI) (De Nettancourt, 1997) or interspecific (e.g. unilateral) incompatibility (UI) (Lewis & Crowe, 1958). An important interspecific RB occurs between closely related species, in which the pollen from one species is rejected by the pistils of another (incompatible response) but not vice versa (unilateral), hence is termed interspecific unilateral incompatibility (UI) (Lewis & Crowe, 1958, Pandey, 1981, Onus & Pickersgill, 2004).

As a major interspecific RB, UI has been documented in many genera (Lewis & Crowe, 1958, Abdalla, 1977, Pandey, 1981, Eijlander, 1998, Wang *et al*., 2022, Huang *et al*., 2023). Studies in the Solanaceae found that UI usually follows the “SI × SC” rule, whereby pollen tubes from a self-compatible (SC) species are rejected in styles of SI species, while in the reciprocal cross, pollen tubes from SI species successfully fertilize ovules of SC species (Lewis & Crowe, 1958, Pandey, 1962, Murfett *et al*., 1996). Thus, in this scenario, many of the molecular mechanisms are thought to be shared between UI and SI systems (Bernacchi & Tanksley, 1997, Li & Chetelat, 2010, Tovar-Méndez *et al*., 2014, Li & Chetelat, 2015). Three stylar UI (*sui*) loci *sui1.1*, *3.1* and *12.1* were identified by quantitative trait loci (QTL) analysis in a BC_1_ population derived from *S. habrochaites* (formerly *Lycopersicon hirsutum*) LA1777 and *S. lycopersicum* (formerly *L. esculentum*) cv. E6203 (Bernacchi & Tanksley, 1997). Two of these (*sui1.1* and *12.1*) were found to be co-localized with the known stylar SI factors *S-RNase* and *HT*, respectively (Bernacchi & Tanksley, 1997, Covey *et al*., 2010, Tovar-Méndez *et al*., 2014, Jewell *et al*., 2020). Genetic and transgenic experiments in *Nicotiana* and *Solanum* demonstrated that S-RNase together with HT proteins produce stylar barriers in both SI and UI systems, although neither alone were sufficient to result in pollen tube rejection (Murfett *et al*., 1994, Murfett *et al*., 1996, McClure *et al*., 1999, Hancock *et al*., 2005, Tovar-Méndez *et al*., 2014). On the pollen side, SLF-23 and CUL1 were demonstrated as pollen factors underlying pollen UI loci *ui1.1* and *6.1*, which are required for breaking the S-RNase-dependent pollen rejection (Chetelat & Deverna, 1991, Li & Chetelat, 2010, Li *et al*., 2010, Li & Chetelat, 2015). However, in contrast to the UI between SI and SC species, there are relatively few studies on the genetic basis of UI between SC species (Murfett *et al*., 1996, Baek, 2011).

UI between SC species is reported in *Solanum* section *Lycopersicon*, which consists of cultivated tomato *S. lycopersicum* and 16 wild tomato species with diverse mating systems (Bedinger *et al*., 2011, Baek *et al*., 2016, Ramírez-Ojeda *et al*., 2021). Recently, the genomes of 10 from 17 tomato species were sequenced (Li *et al*., 2023b). Thus, the tomato clade provides an ideal system with which to study the genetic basis of interspecific UI. Pollen of four SC red-fruited species (*S*. *lycopersicum*, *S*. *pimpinellifolium*, *S*. *cheesmaniae* and *S*. *galapagense*) is rejected by styles of all SI green-fruited species, whereas the reciprocal crosses are compatible (Bedinger *et al*., 2011, Baek *et al*., 2015). Loss of heterospecific pollen rejection in SC red-fruited tomato styles is considered a result of non-functional S-RNase and HT proteins (Kondo *et al*., 2002a, Kondo *et al*., 2002b). However, some SC accessions of wild tomato species which lack of S-RNase activity, e.g. *S. pennellii* LA0716, *S. arcanum* LA2157, *S. habrochaites* LA0407 and *S. chmielewskii* LA1316 (Kondo *et al*., 2002a, Chalivendra *et al*., 2013, Tovar-Méndez *et al*., 2017), still display stylar UI (*sui*) responses to pollen from SC red-fruited species (Covey *et al*., 2010, Bedinger *et al*., 2011). Therefore, sui factor(s) other than S-RNases are supposed to present in these wild tomato species. Abundant transcripts of *HT-A*, one of the two *HT* gene homologs (*HT-A* and *HT-B*), are detected in styles of all examined SC wild tomato accessions, e.g. LA0716, LA0407, LA2860, LA2157 and LA1316 (Kondo *et al*., 2002a, Covey *et al*., 2010, Pease *et al*., 2016, Tovar-Méndez *et al*., 2017). Moreover, the UI response in LA0716 styles is temporally and spatially associated with the accumulation of HT-A proteins (Chalivendra *et al*., 2013). *S. lycopersicum* pollen tubes grew significantly longer in LA0716 transgenic styles with reduced *HT-gene* expression (RNAi) (Tovar-Méndez *et al*., 2017). Moreover, *S. lycopersicum* pollen tubes kept elongating and fertilizing ovules in transgenic styles of LA0407, LA2860 and LA2157 with reduced *HT* expression (Tovar-Méndez *et al*., 2017). These observations suggest that *HT* genes are involved in S-RNase-independent UI mechanisms.

Recently, three UI genes (one *pui* and two *sui* genes) were identified in LA0716 (Qin *et al*., 2018, Muñoz-Sanz *et al*., 2021, Qin & Chetelat, 2021). A forward genetics study demonstrated that a *pui* factor *FPS2*, encoding farnesyl pyrophosphate synthase 2, abundantly expressed in pollen, was identified as *pui10.1* (Chetelat & Deverna, 1991, Qin *et al*., 2018). Given that red-fruited tomato species are self-fertile, *pui* genes are thought to play a specific role in breaking the stylar UI response rather than being requiring for pollen tube elongation in the stylar transmitting tract. Consistent with this hypothesis, pollen tubes of LA0716 with *SpFPS2*^loss-of-function^ (LA0716*^fps2^*) were rejected by LA0716 styles, while they were accepted by styles of *S. lycopersicum* VF36 (Qin *et al*., 2018) (Table S1, black cells). Moreover, pollen tube rejection of LA0716*^fps2^* was observed on styles of IL3-3 as well (Table S1, light grey cells), in which a gene cluster encoding the demonstrated sui factor, ornithine decarboxylases (SpODC2s), was introgressed (Qin & Chetelat, 2021). It is noteworthy that styles of IL3-3 were unable to reject pollen from either *S*. *lycopersicum* (Hamlin *et al*., 2017) or LA0716 (Qin & Chetelat, 2021) (Table S1, light grey cells). When *SpODC2s* were knocked down in styles of either LA0716 or IL3-3, pollen tubes of LA0716*^fps2^* were accepted (Qin & Chetelat, 2021) (Table S1, light grey cells), indicating that pui SpFPS2 and sui SpODC2 are antagonistic in the UI response.

Pollen tubes of either *S. lycopersicum* or LA0716*^fps2^*were rejected in styles of F_1_ (*S. lycopersicum* × LA0716) (Liedl *et al*., 1996, Qin *et al*., 2018) (Table S1, dark grey cells). Interestingly, self-pollinations of F_1_ (*S. lycopersicum* × LA0716*^fps2^*) were compatible resulting in normal seed set (Qin *et al*., 2018) (Table S1, dark grey cells). In other words, compared with parental pollen tubes (*S. lycopersicum* and LA0716*^fps2^*), the offspring pollen tubes of F_1_ (*S. lycopersicum* × LA0716*^fps2^*) were able to break the stylar UI response after segregation and independent assortment of genetic factors. *SlFPS2* in *S. lycopersicum* pollen encodes a protein product with similar amino acid sequence but exhibits significantly lower transcript abundances compared with that of LA0716 pollen (Qin *et al*., 2018). Moreover, in F_2_ (*S. lycopersicum* × LA0716*^fps2^*), the ratio of *SlFPS2* was significantly higher than Mendelian expectations (Qin *et al*., 2018), indicating that *SlFPS2* was preferentially selected compared to the *fps2* allele. These results revealed that in styles of F_1_ (*S. lycopersicum* × LA0716), of the three genotypes of pollen tubes, 1) *SlFPS2* in *S. lycopersicum*, 2) LA0716*^fps2^* and 3) *SlFPS2* with unknown nonallelic loci from LA0716 (contributed by recombination), only pollen tubes of 3) could break the stylar UI response, suggesting that additional pui factor(s) other than SpFPS2 are present in LA0716 (Qin *et al*., 2018).

In this study, pollen from F_1_ (*S. lycopersicum* Heinz × LA0716) was used to backcross styles of LA0716. It is hypothesized that pollen tubes lacking *pui* factor(s) are rejected in styles of LA0716. Therefore, *pui* factors are supposed to exhibit transmission ratio distortion (TRD) in the BC_1_ (Heinz × LA0716). High-throughput sequencing technologies, single nucleotide polymorphism (SNP) genotyping and pollen RNA-seq data were used to identify *pui* loci (Sim *et al*., 2012, Thomson *et al*., 2012). Moreover, introgression lines (ILs) and backcross inbred lines (BILs) derived from LA0716 and *S. lycopersicum* M82 were used for functional validation of *pui* loci (Eshed & Zamir, 1995, Alseekh *et al*., 2013, Hamlin *et al*., 2017). Furthermore, three candidate genes linked to *pui6.2* and *pui12.1* were screened according to our results. In conclusion, we identified that 1) two *pui* QTLs in addition to the reported *pui10.1* (Qin *et al*., 2018), and these three *pui* factors are sufficient to partially overcome the stylar UI response of LA0716; 2) gene candidates underlying the two *pui* loci may be involved in HT-related stylar UI responses; 3) the functional diversity of three *pui* factors in styles of three SC wild tomatoes.

## Materials and methods

### Plant materials

Tomato materials used in the study as shown in (Table S2). These seeds were obtained from the Tomato Genetics Resource Center (https://tgrc.ucdavis.edu/). Tomato plants were cultivated in an artificial climate room (16 h of light at 24 ℃: 8 h of dark at 18 ℃, 50 % humidity).

Progeny of LA0716 by binary mixture pollination was obtained following the method described previously (Guo *et al*., 2019). Mean pollen numbers per flower of LA0716 and Heinz were calculated, and pollen viability was examined (Muhlemann *et al*., 2018). Homogeneous pollen mixtures were prepared by mixing equal amount pollen grains from Heinz and LA0716 and pollinated emasculated flowers of LA0716. 155 seeds were randomly selected for subsequently DNA isolation and genotyping.

### DNA isolation and genotyping

To genotype paternity of seeds produced by pollination of backcrosses and binary pollen mixture, genomic DNA was extracted from ∼50 mg of young leaves of two-week-old seedlings. Extraction was performed using a modified CTAB method (Allen *et al*., 2006). Extracted DNA was treated with RNase A (Bioteke, Cat. No. SP1001) 20 min at room temperature. About 500 ng DNA of each BC_1_ plant was sampled for resequencing. An InDel marker of *CUL1* (Li & Chetelat, 2010) was used to discriminate genotypes between Heinz and LA0716 in the progeny derived from binary mixture pollinations.

*SpODC2* was proven as a stylar UI factor, which was introgressed into IL3-3 (Qin & Chetelat, 2021). *SpHT* was also demonstrated as a S-RNase independent stylar UI factor, which was introgressed into IL12-3 (Bernacchi & Tanksley, 1997, Covey *et al*., 2010, Hamlin *et al*., 2017, Qin & Chetelat, 2021). To discriminate species-specific orthologs of these genes, primers were designed to amplify either *SpODC2c* and *SpHT-A* in LA0716 or *SlODC2* and *SlHT-A* in M82. These primers were also used to examine introgressed segments in IL3-3, IL12-3, BIL6546 and F_2_ (BIL6546 × IL3-3). Meanwhile, the genomic DNA of BIL6546 was sequenced to confirm its introgressed segments from *S. pennellii* (Table S6). Based on genomic sequences of LA0716 and M82, two InDel markers tightly linked to *pui6.2* and *10.1* were used to genotype BIL6546, BIL6676, IL10-1, and related F_2_ populations. All primers used in this study were listed in Table S3.

### DNA resequencing, SNP-calling and QTL analysis

Genomic DNA of BC_1_ population was sequenced using the MGI DNBSEQ T7 platform. Insert sizes of paired-end sequencing libraries were 250–350 base pairs (bp) in length, and read length was 150 bp. For F_1_ and each BC_1_ individual, ∼30 gigabase pair (Gbp) and 10 Gbp of clean data were generated, respectively. BWA (Li & Durbin, 2009) was used for aligning and mapping short reads on the tomato reference genome ITAG4.1 (https://solgenomics.net/organism/Solanumlycopersicum/genome). SNP-calling was performed using SAMtools (Li *et al*., 2009) and GATK toolkit (McKenna *et al*., 2010) with default parameters. For all genome resequencing data, filter parameters in Python scripts were Qual >= 30 and QD > 2.0 and MQ >= 50.0 and FS <= 50.0 and SOR < 3.0 and MQRankSum >= –8 and ReadPosRankSum >= –6. Heterozygous sites of F_1_ progeny were extracted with a genotype of 0/1 (0.3 <= the SNP-index <= 0.7, 0: genotype of Heinz (L), 1: LA0716 (P)) and used to define the SNPs distinguishing the two parents (Heinz and LA0716). For the BC_1_ population, sites with genotypes of 0/1 or 1/1 were extracted by screening a subset of SNP markers (missing data < 10% and F_1_ heterozygous sites). To calculate the average DPG_P–L_ of a gene, the SNPs located in the regions, 1) 2 Kbp (kilobase pair) upstream of the gene, 2) gene body, and 3) 1 Kbp downstream of the gene, were used.

### Statistical analysis

Pollen tube length and RT-qPCR data were visualized using GraphPad Prism 9 software. Statistical differences of pollen tube growth among different genotypes were calculated by one-way ANOVA with a Scheffe test using IBM SPSS Statistics 27 software. The significance of observed and expected genotypic frequencies of F_2_ (BIL6546 × IL3-3) offspring was evaluated using binomial distribution test. Meanwhile, Chi-square goodness-of-fit statistics were calculated to assess the significance of transmission ratio distortion (TRD) at *ODC2*, *HT-A* and *pui6.2* (Qin & Chetelat, 2021).

**Other methods corresponding to the supplemental data could be found in supplemental information**.

## Results

### Two pollen UI QTLs, *pui10.1* and *pui6.2*, were characterized by resequencing BC_1_ population derived from *S. lycopersicum* Heinz and *S. pennellii* LA0716

*SpFPS2* was previously characterized as a *pui* factor in LA0716 (Qin *et al*., 2018). As mentioned above, *SlFPS2* with additional *pui* factor(s) could break the stylar UI response of F_1_ (*S. lycopersicum* × LA0716) (Qin *et al*., 2018) (Table S1, dark grey cells). To mine *pui* factors in LA0716, a BC_1_ population was constructed between Heinz (LA4345) and LA0716 (Fig. 1a) as follows: 1) styles of Heinz were pollinated by pollen grains from LA0716 to generate F_1_ (Heinz × LA0716); 2) pollen from F_1_ (Heinz × LA0716) were used to pollinate styles of LA0716 to create the BC_1_ population. After segregation and independent assortment, pollen tubes of F_1_ (Heinz × LA0716) with *pui* factor(s) penetrate styles of LA0716 and fulfill the double fertilization required for seed set, while those without *pui* factor(s) are rejected by styles and fail to fertilize ovules. In other words, in the BC_1_ population, paternal genomic regions containing *pui* QTL(s) are supposed to deviate from Mendelian expectations and exhibit transmission ratio distortions (TRDs). Considering compatible and incompatible pollen grains are mixed in pollen grains of F_1_ (Heinz × LA0716), it is not clear whether the pollen mentor effect (Lan *et al*., 2023) would be present in this backcross. To answer this question, we used equal pollen mixtures from Heinz and LA0716 to pollinate LA0716 styles. No Heinz genotype was observed in 155 offspring by the InDel marker of *CUL1* (Li & Chetelat, 2010) (Fig. S1), suggesting that pollen mentoring is not occurring in these crosses, which is consistent with a past report of UI between *S. lycopersicum* cv. New Yorker and SC S. pennellii LA2963 (Liedl *et al*., 1996).

**Fig. 1.**
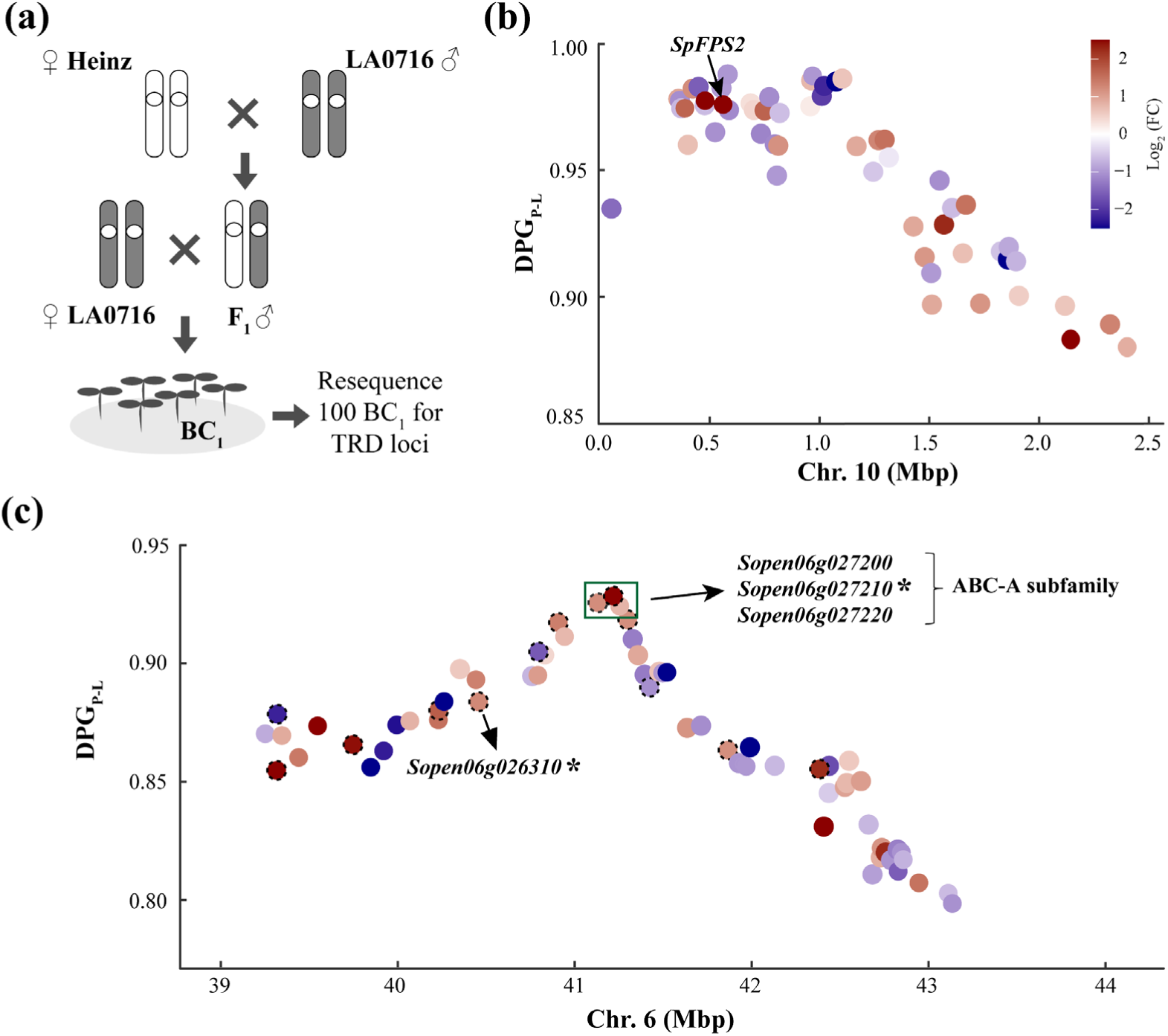
Identification of *pui* loci by resequencing 100 BC_1_ individuals between *S. lycopersicum* Heinz and *S. pennellii* LA0716. (a) Schematic diagram of BC_1_ population construction between Heinz and LA0716. In (b) and (c), each dot represents gene with significant difference (*P* < 0.05) in expression in pollen between LA0716 and Heinz (transcriptome data). Differences of paternal genotype (DPG) ratios between *pennellii* (P) and *lycopersicum* (L) tomato (DPG_P – L_, Y axis) of genes on Chr. 10 (b) and 6 (c) are shown, as well as chromosomal location (X axis, million base pair, Mbp) and dot color (transcript abundance heatmap of fold change (FC) in pollen between LA0716 and Heinz). Dot corresponding to the known *SpFPS2* at *pui10.1* is indicated by black arrow in (b). In (c), dots corresponding to gene candidates with more than 2-fold differences in transcript abundances between LA0716 and Heinz pollen are circled in black dash line and chosen for further functional analyses. In *pui6.2*, three ABC-A transporter homologous genes were linked to DPG_P – L_ peak and labeled by a green box. Asterisks indicate genes whose protein product could interact with the HT protein in yeast (Fig. S5b and S6b).

To identify the paternal locus of TRD, 100 BC_1_ individuals were randomly chosen for resequencing. About 12.55 million SNPs were used to analyze the resequencing data. In the BC_1_ population, as to the non-*pui* locus, the difference of parental genotype ratio between *pennellii* (P) and *lycopersicum* (L) tomatoes (DPG_P–L_) is expected to be 0.5 (75%–25%). The DPG_P–L_ of a *pui* QTL should be significantly higher than 0.5. Two TRD loci were detected: one locus is located on chromosome (Chr.) 10 (DPG_P–L_ = 0.99, *χ^2^* = 96.04; *χ^2^*_1e-21 = 91.72, *P* < 1e-21), which is considered as *pui10.1* (Fig. 1b); another one is located on Chr. 6 (DPG_P–L_ = 0.93, *χ^2^* = 73.96; *χ^2^*_1e-17 = 73.51, *P* < 1e-17) (Fig. 1c). Since *ui6.1* encodes *CUL1* involved in UI between SI and SC tomatoes (Li & Chetelat, 2010, Li *et al*., 2010), the *pui* QTL at Chr. 6 in this study was named as *pui6.2*.

Prior studies suggested that genes contributing to UI exhibited functional differences (e.g. gene structural variations or different transcript abundance) between UI– and non-UI-competent styles among different tomato species, e.g. *HT*-*A*/*B* and *ODC2s* (Bedinger *et al*., 2011, Chalivendra *et al*., 2013, Pease *et al*., 2016). Also, reported *pui* factor *FPS2* transcribes more in LA0716 pollen *versus S. lycopersicum* pollen, and acts in a dose-dependent manner (Qin *et al*., 2018, Qin & Chetelat, 2021). Therefore, a gene encoding a pui factor is hypothesized to be: 1) located at a locus with significantly higher TRD compared to other loci; 2) displaying functional difference in pollen between Heinz and LA0716. Thus, we used pollen transcriptome data of LA0716 and Heinz to filter genes tightly linked to the *pui* QTL. Gene filtering requirements were: 1) annotated genes located at a region tightly linked to the *pui* QTL: 337 genes located at the region of 0 to 2.5 Mbp at Chr. 10 and 577 genes 39.2 to 43.2 Mbp at Chr. 6; 2) based on pollen transcriptome data of both LA0716 and Heinz (TPM > 1 in both transcriptomes), 130 genes linked to Chr. 10 and 194 genes linked to Chr. 6 were further selected; 3) differentially expressed genes (DEGs, n = 3, *P*< 0.05): 55 pollen DEGs linked to *pui10.1* (Fig. 1b and Table S4) and 61 pollen DEGs linked to *pui6.2* (Fig. 1c and Table S5) were identified. Consistent with the reported *ui10.1* (Qin *et al*., 2018), the *pui* factor *SpFPS2* (DPG_P–L_ = 0.9758 in this study) was found among these 55 genes linked to *pui10.1* (Fig. 1b). Therefore, we inferred that *SpFPS2* was the gene underlying *pui10.1*.

Further analysis was performed to characterize *pui6.2*. Candidate genes showing either higher or lower transcript abundance in LA0716 pollen *versus* Heinz were selected. Among 61 pollen DEGs (Table S5), 13 genes showed more than 2-fold differences in transcript abundance between LA0716 and Heinz (Table S6) and were selected as candidate genes for further functional validation.

### Functional analysis of *pui6.2*

To determine whether *pui6.2* is involved in the pollen UI response, available ILs and BILs that *S. lycopersicum* M82 (LA3475) with homozygous introgressed chromosomal segments from *S. pennellii* LA0716 (Eshed & Zamir, 1995, Sim *et al*., 2012, Ofner *et al*., 2016) were used in this study. In pollen of IL10-1 (with introgressed *pui10.1*, *SpFPS2*) (Fig. S2a and Table S7), RT-qPCR assays revealed ∼11 times higher transcript abundance of *SpFPS2* compared with M82 (Fig. S2b). The *pui6.2* was introgressed into IL6-2 and BIL6676 as shown in schematic diagram (Fig. S3) (Eshed & Zamir, 1995, Sim *et al*., 2012, Ofner *et al*., 2016). Due to the described strong necrotic phenotype of IL6-2 (https://tgrc.ucdavis.edu/pennelliiILs), we used BIL6676 to verify the function of *pui6.2*. Pollen containing either 1) single *pui10.1* (IL10-1), 2) single *pui6.2* (BIL6676) or 3) dual *pui10.1 pui6.2* (F_1_ (IL10-1 × BIL6676)) were used to pollinate LA0716 styles. Like M82, pollen tubes of all three genotypes were rejected in the upper part in LA0716 styles 72 h after pollination (HAP) (Fig. 4a, green panel and Fig. S2c), suggesting that the function of *pui10.1* and *pui6.2* in IL/BIL might be too weak to break the stylar UI response of LA0716.

We hope that a weakened stylar UI system derived from M82 and LA0716 is suitable for the function validation of *pui* factors. The reported *sui* factor *SpHT-A*/*B* can be found in the corresponding IL (IL12-3) and BIL (BIL6546) materials (Fig. S4c, S4e, S4f and Table S7). In *SlHT-A*, there is an insertion of 14 base pairs at the second exon compared to *SpHT-A* (Fig. S4a), leading to a frame shift, a premature stop codon and the loss-of-C terminus (Fig. S4b), suggesting that SlHT-A has lost its function. In *SlHT-B*, only trace amounts of transcripts were found in styles of M82 compared with others (Fig. S4f), implying functional variation of HT-B between M82 and LA0716. In the styles of IL12-3 and BIL6546, the transcript abundance of either *HT-A* or *HT-B* is significantly lower than that of LA0716 (Fig. S4e and S4f), suggesting they could be used to generate weakened stylar UI systems.

Another reported *sui* factor is *SpODC2s*, whose amino acid sequences are similar between M82 and LA0716, but expressed 1000-fold higher in LA0716 styles compared with M82 styles (Qin & Chetelat, 2021) (Fig. S4d). The IL corresponding to *SpODC2s* is IL3-3 (Fig. S4c and Table S7). In styles of IL3-3, the transcript abundance of *SpODC2s* is similar to that of LA0716 and significantly higher than that of M82 (Fig. S4d).

Recently, styles of F_1_ (IL12-3 ×IL3-3) were demonstrated to reject pollen tubes of *S. lycopersicum* (Hamlin *et al*., 2017, Qin & Chetelat, 2021). Thus, we created two hybrids: F_1_ (IL12-3 × IL3-3) as described in (Hamlin *et al*., 2017, Qin & Chetelat, 2021)(Fig. 2a) and F_1_ (BIL6546 × IL3-3) (Fig. 2c and S4d-f) to validate the function of *pui6.2*. Compared with styles of LA0716, both M82 and Heinz pollen tubes grew significantly longer and were finally rejected in styles of either F_1_ (IL12-3 × IL3-3) or (BIL6546 × IL3-3) (Fig. 2b and 2d), suggesting that styles of the two F_1_ hybrids containing both SpHT-A/B and SpODC2 are weakened stylar UI systems. It is noteworthy that BIL6546 contains not only *SpHT-A*/*B* but also *pui6.2* (Fig. S4 and Table S7). In styles of the two hybrids, pollen tubes containing either *pui10.1* or *pui6.2* could break the stylar UI response and result in fruit and seed set (Fig. 2b and 2d). Compared with the size of the *pui6.2* locus (Fig. 1c, ∼40 to 42 Mbp, peak of TRD is at ∼41.23 Mbp), introgression of the fragment of Chr. 6 in either BIL6676 (∼39.28 to 45.95 Mbp, containing 923 genes) or BIL6546 (Table S7, ∼0.06 to 35.73 Mbp, containing 1204 genes and ∼40.39 to 42.5 Mbp, containing 279 genes) is too large to provide a validation of *pui6.2* function. Clearly, more precise evidence is needed.

**Fig. 2.**
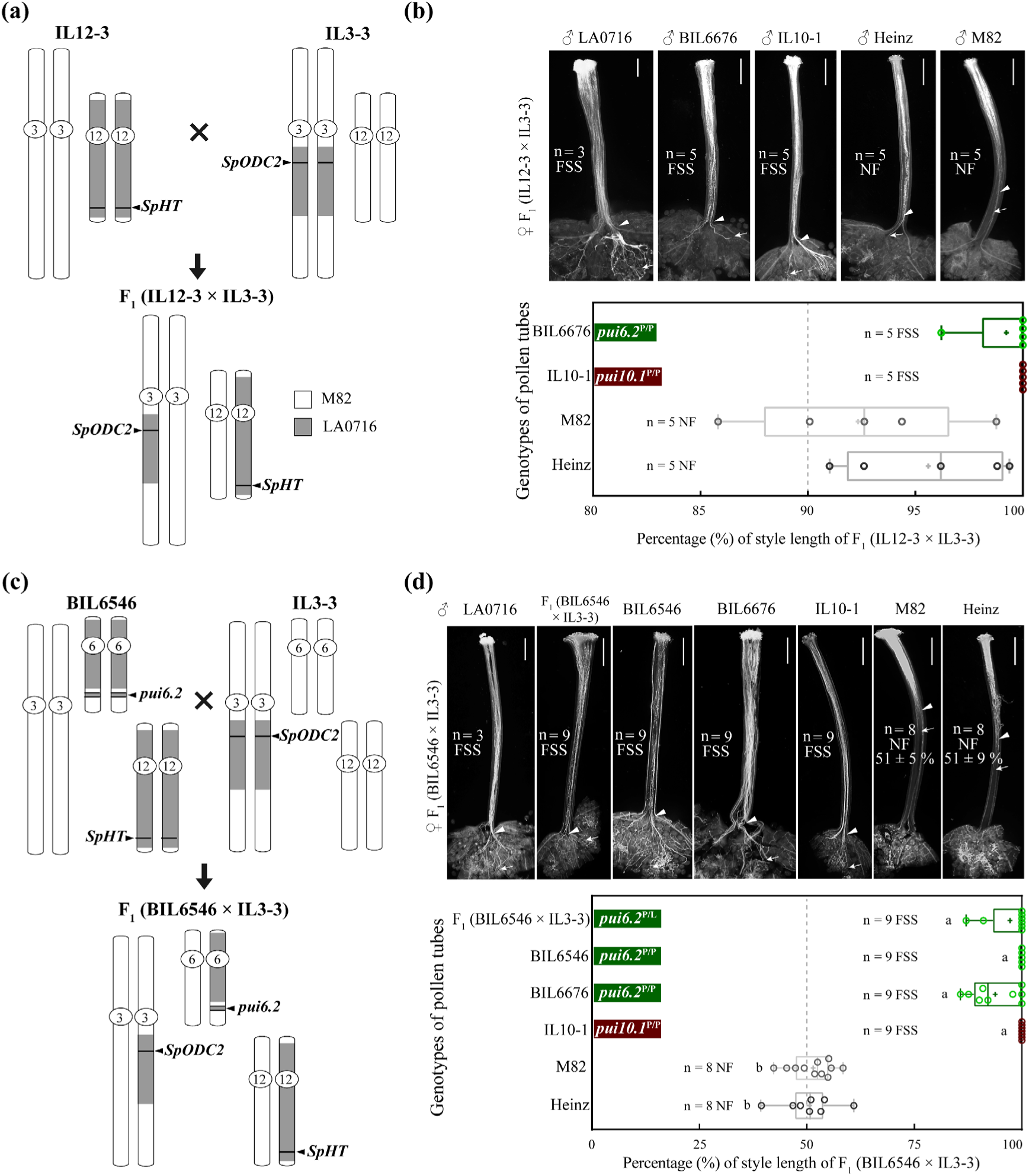
Cultivated tomato pollen with introgressed *pui6.2* breaks the stylar UI response resulting from *sui3.1* (*SpODC2*) and *sui12.1* (*SpHT*). (a) Genotype schematic diagram of F_1_ (IL12-3 × IL3-3) is shown. Introgressed fragments from LA0716 (shown in grey) are located on Chr. 3 and 12 (other chromosomes are from M82). (b) Pollen tube staining by aniline blue (upper panel) and length measurement (bottom panel) in styles of F_1_ (IL12-3 × IL3-3) 48 h after pollination. Genotypes of pollen tubes as indicated: IL10-1 contains introgressed *SpFPS2* (on *pui10.1*) (Qin *et al*., 2018); BIL6676 contains above-mentioned *pui6.2*. (c) Genotype schematic diagram of F_1_ (BIL6546 × IL3-3) was shown, whose introgressed fragments from LA0716 (shown in grey) are located on Chr. 3, 6 and 12 (other chromosomes are from M82). Introgressed segments were determined by SNP markers according to (Sim *et al*., 2012), and validated by genomic DNA resequencing of IL3-3 and BIL6546 (Table S7). (d) Pollen tube staining by aniline blue (upper panel) and length measurement (bottom panel) in styles of F_1_ (BIL6546 × IL3-3) 48 h after pollination. Genotypes of pollen tubes as indicated: IL10-1 contains introgressed *SpFPS2* (on *pui10.1*) (Qin *et al*., 2018); BIL6676, BIL6546 and F_1_ (BIL6546 × IL3-3) contain above-mentioned *pui6.2*. For (a) and (c), chromosomal segments of M82 are shown in white and a scale representing physical length of the chromosomes is shown. For (b) and (d), white triangles and white arrows represent a majority of pollen tubes and the longest pollen tubes in pistils, respectively. FSS: fruit and seed set; NF: no fruit. Scale bar, 1 mm. Length of pollen tubes were represented by circles (each circle corresponds a pollinated style as a biological replicate) in the box-and-whisker plot, + indicates mean value of data. Different letters indicated significant differences (one-way ANOVA (Scheffe test), *P* < 0.05). Green and red rectangle represent introgressed *pui6.2* and *10.1* QTLs, respectively. P/P and P/L in superscript indicate homozygous and heterozygous *pui* locus, respectively.

In F_1_ (BIL6546 × IL3-3), the two *sui* factors (*SpODC2* and *SpHT-A/B*) as well as *pui6.2* are heterozygous. Given that F_1_ (BIL6546 × IL3-3) could break the stylar UI response of F_1_ (BIL6546 × IL3-3) and produce seeds (Fig. 2d), we inferred that in F_2_ (BIL6546 × IL3-3), the transmission rate of *pui6.2* (LA0716 allele) should be significantly higher than Mendelian expectation. To test this hypothesis, we analyzed the genotypes of *pui6.2* (genotyping primer pairs of *pui6.2*-IDL-F and –R are listed in (Table S3), and their PCR product is at ∼41.23 Mbp on Chr. 6) in 194 F_2_ (BIL6546 × IL3-3) individuals (Fig. 3a). Consistent with our hypothesis, an extreme TRD on *pui6.2* was observed with a 9 LL: 103 PL: 82 PP genotype ratio (*χ^2^* = 55.68, *P* = 2.03E-41) as shown in (Table 1).

**Fig. 3.**
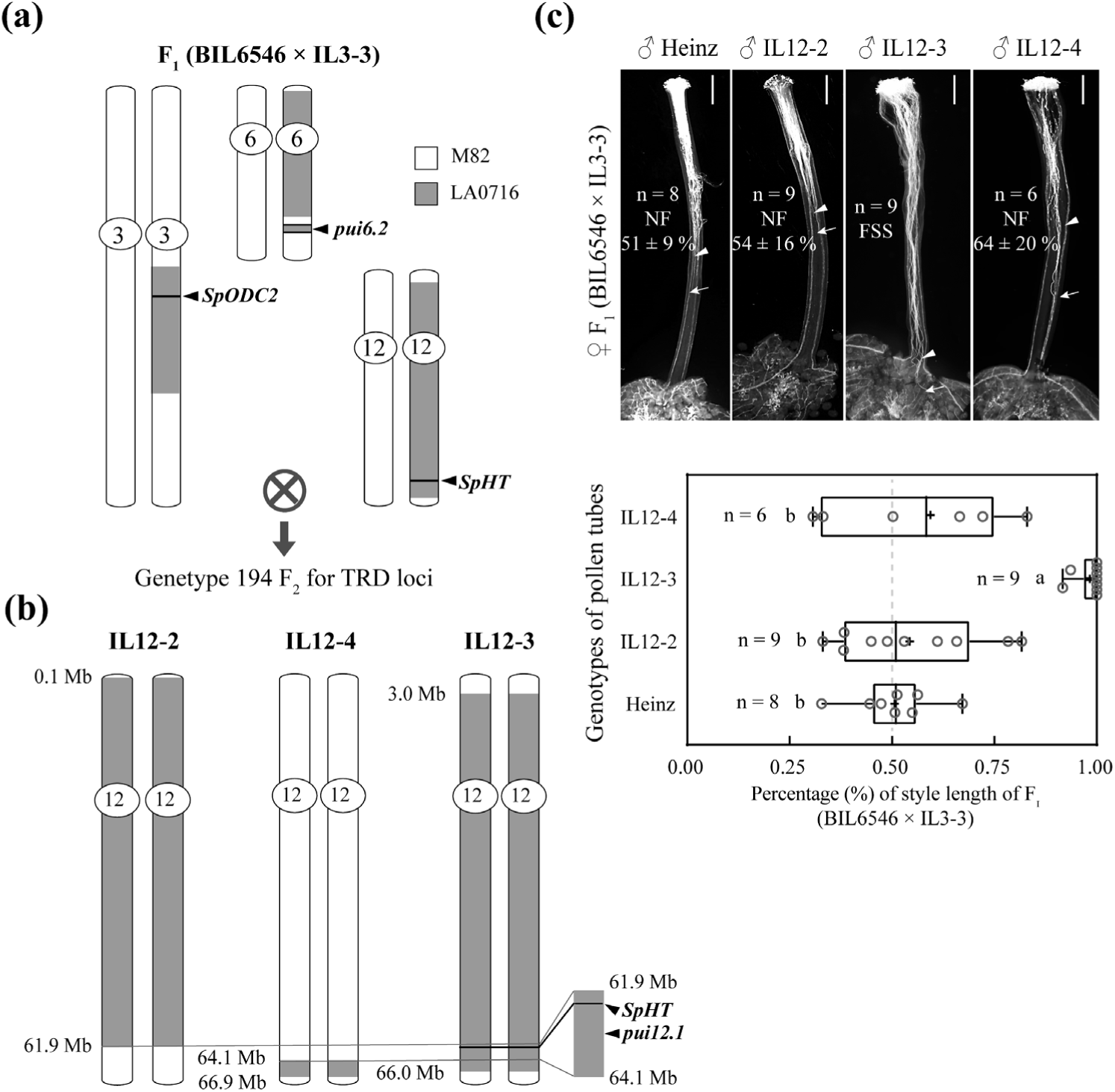
Pollen from IL12-3 with introgressed *pui12.1*, but not IL12-2 and 12-4, could break the UI response in F_1_ (BIL6546 × IL3-3) styles and accomplish fertilization. (a) Schematic diagram of experimental design to determine if *pui* loci are linked to introgressed fragments on Chr. 3, 6 and 12. (b) Chromosome schematic diagram of IL12-2, 12-3 and 12-4. The three ILs all have an introgressed segment from *S. pennellii* LA0716 on Chr. 12 (as shown in grey). According to SNP markers, the region from 61.9 to 64.1 Mbp is unique to IL12-3 (grey rectangle on the right), named *pui12.1*, and specifically contains the stylar UI factors *SpHT-A*/*B*. (c) Upper panel: pollen tube staining by aniline blue on styles of F_1_ (BIL6546 × IL3-3) 48 h after pollination. FSS: fruit and seed set. NF: no fruit. White triangles and arrows represent a majority of pollen tubes and the longest pollen tubes in pistils, respectively. Scale bar, 1 mm. Bottom panel: length of pollen tube growth is represented by circles (each circle corresponds a pollinated style as a biological replicate) in the box-and-whisker plot, + indicates mean of data. Different letters indicated significant differences (one-way ANOVA (Scheffe test), *P* < 0.05).

**Table 1.** Transmission ratio distortion (TRD) occurred at two of three heterozygous loci in F_2_ (BIL6546 × IL3-3), which are namely *pui6.2* and *12.1*.

| Chr.3_*ODC2* | Chr.12_*HT-A* |  |  | Chr.6_*pui6.2* |
| --- | --- | --- | --- | --- |
|  | LL | PL | PP |  |
| LL | 0 (3.03) * | 0 (6.06) ** | 2 (3.03) | LL |
|  | 2 (6.06) * | 12 (12.12) | 12 (6.06) * | PL |
|  | 2 (6.06) | 12 (6.06) * | 7 (3.03) * | PP |
| PL | 0 (6.06) ** | 2 (12.12) *** | 2 (6.06) * | LL |
|  | 4 (12.12) ** | 22 (24.24) | 22 (12.12) ** | PL |
|  | 3 (6.06) | 14 (12.12) | 22 (6.06) *** | PP |
| PP | 0 (3.03) * | 1 (6.06) * | 1 (3.03) | LL |
|  | 0 (6.06) ** | 20 (12.12) ** | 8 (6.06) | PL |
|  | 1 (6.06) | 10 (6.06) * | 12 (3.03) *** | PP |

| Chr.3_*ODC2* | LL | PL | PP | Chr.12_*HT-A* | LL | PL | PP |
| --- | --- | --- | --- | --- | --- | --- | --- |
| Expected | 48.5 | 97 | 48.5 | Expected | 48.5 | 97 | 48.5 |
| Observed | 49 | 92 | 53 | Observed | 12 | 93 | 89 |
| $X^2_{\text{Chr.3\_ODC2}}$ | 0.68 | $P = 0.7193$ | | $X^2_{\text{Chr.12\_HT-A}}$ | 61.45 | $P = 7.13\text{E-}29$ | |

| Chr.6_*pui6.2* | LL | PL | PP |
| --- | --- | --- | --- |
| Expected | 48.5 | 97 | 48.5 |
| Observed | 9 | 103 | 82 |
| $X^2_{\text{Chr.6\_pui6.2}}$ | 55.68 | $P = 2.03\text{E-}41$ | |

LL, homozygous *S. lycopersicum* genotype; LP, heterozygous genotype; PP, homozygous *S. pennellii* genotype. Expected values are in parentheses. The observed genotype, whose transmission ratio represented a significant distortion from expected, was indicated by \* ( $P < 0.05$ ), \*\* ( $P < 0.01$ ) or \*\*\* ( $P < 0.001$ ). Exact $P$ -values of genotypes are shown in (Table S8).

Taken together, these results indicated that cultivated tomato pollen tubes with introgressed *pui6.2* could break the stylar UI response contributed by *SpODC2* and *SpHT-A*/*B* (Hamlin *et al*., 2017, Qin & Chetelat, 2021), indicating that *pui6.2* plays a role in UI between Heinz and LA0716.

### Candidate gene prediction on *pui6.2*

*pui* factors are thought to exhibit functional differences between cultivated and *pennellii* tomatoes. To identify candidate gene(s) underlying *pui6.2*, we selected 13 genes (Table S6) to examine their transcriptional abundance in pollen of F_1_ (BIL6546 × IL3-3), M82 and LA0716. As shown in (Fig. S5a), transcripts of 12 genes were detected (except *Sopen06g028230*) in pollen. Within the 12 genes, significant transcriptional differences of 6 genes were observed between F_1_ (BIL6546 × IL3-3) and M82 pollen (*P*< 0.05). The pui6.2 factor was demonstrated to break the stylar UI response established by SpODC2 and SpHT proteins in F_1_ (IL12-3 × IL3-3) and (BIL6546 × IL3-3) (Fig. 2b and 2d), suggesting that pui6.2 associates with either SpODC2s or SpHT proteins.

SpODC2, as an ornithine decarboxylase, is known to antagonize SpFPS2 specifically (Table S1, light grey cells) and probably participates in UI via its enzymatic activity (decarboxylating ornithine to putrescine) (Qin & Chetelat, 2021). No signal peptide is predicted from the ODC2 amino acid sequence (https://services.healthtech.dtu.dk/services/SignalP-6.0/), implying that ODC2 accomplishes its function within stylar cells. HT protein is a small asparagine-rich peptide secreted into the apoplast from stigmatic and transmitting tract cells (McClure *et al*., 1999). Therefore, the protein product encoded by *pui6.2* is hypothesized to directly interact with SpHT. Considering that there are transmembrane domains predicted from the candidate proteins, a mating-based split-ubiquitin (mbSUS) yeast two hybrid (Y2H) assay was used to examine interaction between 6 protein candidates with SpHT-A. Y2H results (Fig. S5b) revealed that two proteins, an ABC-A transporter (encoded by *Sopen06g027210*) and a eukaryotic peptide chain release factor GTP-binding subunit eEF1A (encoded by *Sopen06g026310*), directly interacted with SpHT-A in yeast. Taken together, *Sopen06g027210* and *Sopen06g026310* are screened as candidate genes for *pui* factors underlying *pui6.2* and require further validation.

### Identification of *pui12.1* and candidate gene prediction

Previous studies reported the existence of two additional *pui* factors, one was linked to the introgressed *SpODC2*s (∼2 Mbp away from *SpODC2*s) (Qin & Chetelat, 2021) and another in the region of overlap between IL12-2 and IL12-3 (5.27 to 61.8 Mbp on Chr. 12), but not IL12-3-1 and IL12-4 (Hamlin *et al*., 2017, Qin & Chetelat, 2021). In this study, no peak of TRD was found on either Chr. 3 or Chr. 12 in BC_1_ (Heinz × LA0716). To further validate the potential *pui* factor(s) located on Chr. 3 and Chr. 12, primer pairs to distinguish either *SpODC2 versus SlODC2* or *SpHT-A versus SlHT-A* (Fig. S4c) were used to genotype 194 F_2_ (BIL6546 × IL3-3) individuals (Fig. 3a). Genotyping results demonstrated that the genotype ratio of *ODC2* was close to Mendelian segregation ratio (Table 1, 49 LL: 92 PL: 53 PP, *χ^2^*= 0.68, *P* = 0.7193). However, the genotype ratio of *HT-A* was 12 LL: 93 PL: 89 PP, which significantly deviated from Mendelian expectations (*χ^2^*= 61.45, *P* = 7.13E-29) (Table 1), suggesting a *pui* factor (Bernacchi & Tanksley, 1997, Hamlin *et al*., 2017) (herein *pui12.1*) linked to *SpHT-A/B*. Furthermore, in the 194 examined F_2_ plants, we found none with homozygous LL genotypes on both *pui6.2* and *pui12.1* (*χ^2^* = 12.12, *P* = 2.33E-03), as well as plants with homozygous PP genotypes on both *pui6.2* and *12.1* were significantly higher than Mendelian expectations (41 observed vs 12.12 expected, *χ^2^* = 73.68, *P* = 9.99E-17) (Table 1). In summary, our data suggest that a *pui* locus on Chr. 12 (*pui12.1*) is tightly linked to *SpHT-A*.

Since TRD as extreme as *pui10.1* and *6.2* was not observed on Chr. 12 in BC_1_ (Heinz × LA0716), further analysis is required to confirm the role of *pui12.1*. We used three ILs: IL12-2, IL12-3, and IL12-4 (Fig. 3b) as pollen donors to pollinate styles of F_1_ (BIL6546 × IL3-3). The pollen tube staining assay revealed that pollen tubes from IL12-3, rather than the other two ILs were able to break the stylar UI response of F_1_ (BIL6546 × IL3-3) and fertilize ovules (Fig. 3c), suggesting that *pui12.1* was located at the exclusive region of IL12-3 in comparison with IL12-2 and 12-4 (Fig. 3b). Based on molecular markers reported previously (Eshed & Zamir, 1995, Sim *et al*., 2012), on Chr. 12, introgressed segment in IL12-2 spans approximately 0.1–61.9 Mbp, IL12-3 3–66 Mbp and IL12-4 64.1–66.9 Mbp (Fig. 3b and Table S6). Therefore, the region from 61.9 Mbp to 64.1 Mbp, which is unique to IL12-3, was designated as *pui12.1*. Genes in *pui12.1* were filtered as mentioned above and 12 candidate genes were identified (Table S9). Among these genes, 6 showed at least a 2-fold higher transcript abundance in LA0716 pollen compared with that of Heinz, while the other showed at least 2-fold lower abundances. As mentioned before, only a relatively weak distortion was detected at this region (DPG_P–L_ ranging from 0.6 to 0.79, *χ^2^*: 4–25, *P*-value: 0.045–5.73E-07), implying that other *pui* QTL(s) might function redundantly with *pui12.1*.

In 12 candidate genes, 5 in pollen of F_1_ (BIL6546 × IL3-3) exhibit higher transcript abundances compared with M82 (Fig. S6a and Table S9). Results of Y2H assays revealed that the WD repeat (WDR) protein encoded by *Sopen12g029440* interact with SpHT-A (Fig. S6b). Meanwhile, another homologous WDR protein encoded by *Sopen04g028290* (∼ 66.8% of amino acid identity with Sopen12g029440) (Fig. S6c) could interact with SpHT-A as well (Fig. S6b). The genotype frequency of *Sopen12g029440* in BC_1_ (Heinz × LA0716) was analyzed: 77 PP: 23 PL was observed in the BC_1_ population, which significantly deviated from the expected ratio of 50 PP: 50 PL (*χ^2^* = 29.16, *P* = 1.40E-10) (Table S10, black cells). Moreover, when genotype frequencies were analyzed for the two genes (*Sopen12g029440* and *Sopen04g028290*) simultaneously, only 8 BC_1_ individuals showed PL genotypes on both two genes (deviated from the expected ratio of 25 PL, *χ^2^* = 33.52, *P* = 1.65E-11) (Table S10, grey cell). Our data indicate that the pair of WDR proteins may redundantly function as pui factors.

### *pui6.2*, *10.1* and *12.1* act synergistically to break the stylar UI response of LA0716

Pollen tubes of M82 were rejected in styles of LA0716 (growth arrest occurred at ∼27 % of the style length 72 HAP) (Fig. 4a, green panel). To test if the three introgressed *pui* loci are able to break the stylar UI response of LA0716, 10 genotypes of pollen were used to pollinate LA0716 styles (Fig. 4a), including 1) wild type of M82 and LA0716; 2) three genotypes with single *pui* factors: BIL6676 (containing *pui6.2*), IL10-1 (containing *pui10.1*) and IL12-3 (containing *pui12.1*); 3) three genotypes of pollen with two *pui* factors: BIL6546 (containing both *pui6.2* and *12.1*), F_1_ (IL10-1 × BIL6676) containing both *pui6.2* and *10.1*, and F_1_ (IL10-1 × IL12-3) containing both *pui10.1* and *12.1*; 4) two genotypes containing three *pui* factors: F_1_ and F_2_ (IL10-1 × BIL6546) (note that the three *pui* loci are heterozygous in F_1_ and homozygous in F_2_ (IL10-1 × BIL6546)). Compared with other genotypes of pollen tubes (except LA0716), pollen tubes of F_1_ and F_2_ (IL10-1 × BIL6546) grew significantly longer: F_1_ reached ∼62 % of the LA0716 style 72 HAP, and F_2_ (IL10-1 × BIL6546) ∼87 % (Fig. 4a). It is worth mentioning that in 4 of 17 pollinated LA0716 styles, pollen tubes of F_2_ (IL10-1 × BIL6546) arrived at bottom of the style and resulted in fruits (Fig. 4b). Taken together, these results indicate that a combination of *pui* factors underlying *pui6.2*, *10.1* and *12.1* is sufficient to partially overcome the stylar UI response of LA0716.

**Fig. 4.**
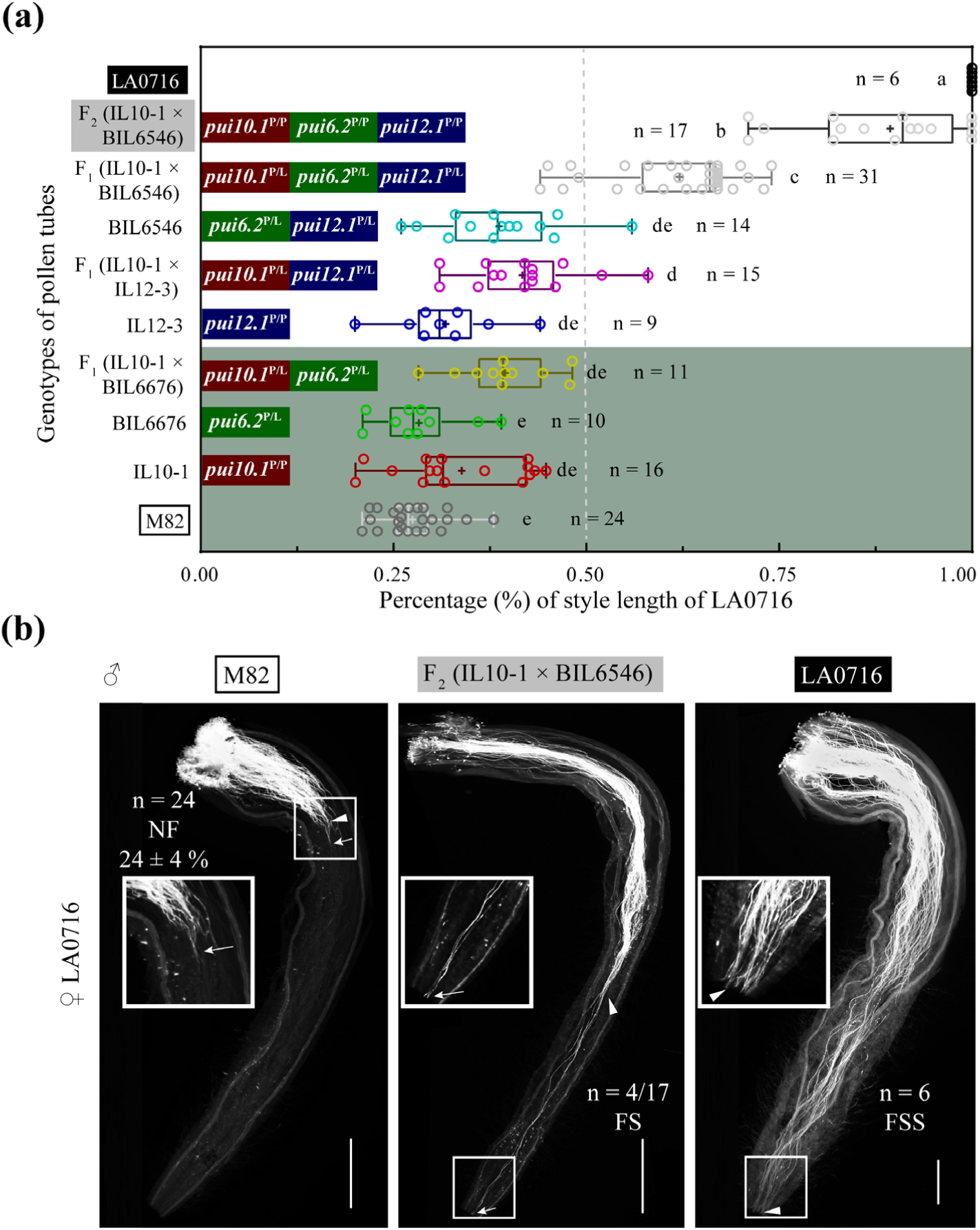
Cultivated tomato pollen tubes with three introgressed *pui* QTLs break the stylar UI response in LA0716 styles. (a) Box-and-whisker plot of pollen tube growth on LA0716 styles 72 h after pollination. Genotypes of pollen tubes as indicated: IL10-1 contains introgressed *SpFPS2* (at *pui10.1*) (Qin *et al*., 2018); BIL6676 contains *pui6.2* from panel A; F_1_ (IL10-1 × BIL6676) contains both *pui10.1* and *6.2*; IL12-3 contains *pui12.1* from panel A; F_1_ (IL10-1 × IL12-3) contains both *pui10.1* and *12.1*; BIL6546 contains both *pui6.2* and *12.1*; F_1_ and F_2_ (IL10-1 × BIL6546) contain all three *pui* loci *pui10.1*, *6.2* and *12.1*. Note that the three *pui* loci are heterozygous in F_1_ (IL10-1 × BIL6546) and homozygous in F_2_. Length of pollen tube growth is represented by circles (each circle corresponds a pollinated style as a biological replicate) in the plot, + indicates mean value of data. Different letters indicated significant differences (one-way ANOVA (Scheffe test), *P* < 0.05). Green, red and blue rectangles represent introgressed *pui6.2*, *10.1* and *12.1*, respectively. P/P and P/L in superscript indicate homozygous and heterozygous *pui* locus, respectively. Data shown in bottom green panel corresponds to section 2 in Results (functional analysis of *pui6.2*). (b) Pollen tube staining by aniline blue in styles of LA0716 72 h after pollination. M82 represents 24 LA0716 styles pollinated by M82 pollen; F_2_ (IL10-1 × BIL6546) represents 4 of 17 LA0716 styles pollinated by F_2_ (IL10-1 × BIL6546) pollen; LA0716 represents 6 styles after self-pollination. NF: no fruit; FS: fruit set; FSS: fruit and seed set. White triangles and arrows represent a majority of pollen tubes and the longest pollen tubes in styles, respectively. Scale bar, 1 mm.

### *pui12.1* is involved in breaking the stylar UI responses of *S. habrochaites* LA0407 and *S. chmielewskii* LA1028

In styles of another two SC wild tomatoes LA0407 and LA1028, LA0716 pollen tubes were accepted (Fig. 5a and 5b), whereas M82 pollen tubes were rejected. 72 HAP, M82 pollen tubes stopped growing at either ∼75 % of the LA0407 style length (Fig. 5a) or ∼83 % of the LA1028 style length (Fig. 5b). To test whether *pui6.2*, *10.1*, and *12.1* from LA0716 function conservatively in breaking stylar UI responses of LA0407 and LA1028, we used the above-mentioned genotypes of pollen for pollination and pollen tube staining, respectively.

**Fig. 5.**
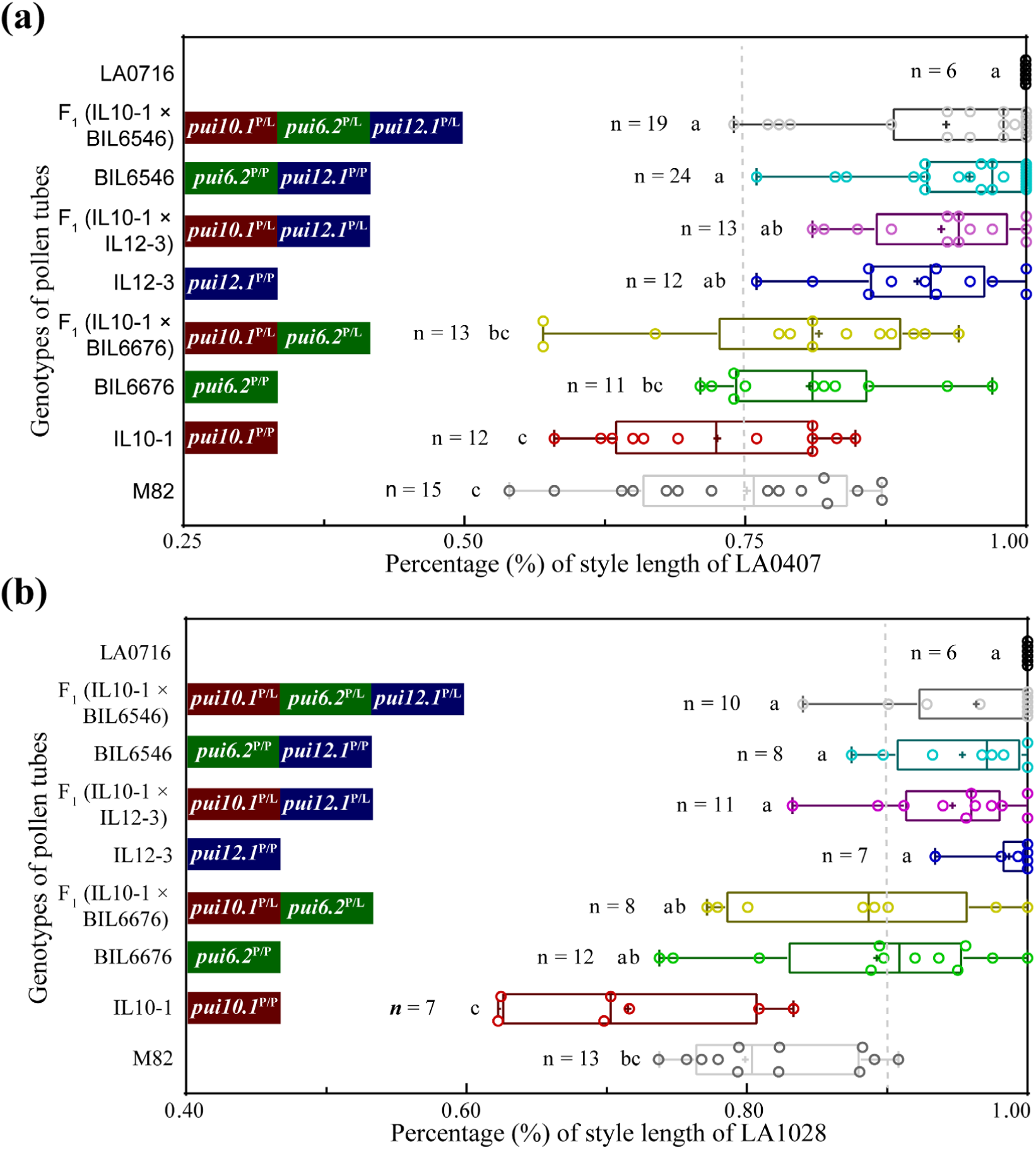
p*u*i12*.1* from LA0716 breaks the stylar UI response between cultivated and another two SC wild tomato species. Box-and-whisker plot of pollen tube growth on styles of *S. habrochaites* LA0407 (a) and *S. chmielewskii* LA1028 (b) 72 h after pollination. Genotypes of pollen tubes as indicated: IL10-1 contains introgressed *SpFPS2* (on *pui10.1*); BIL6676 contains above-mentioned *pui6.2*; F_1_ (IL10-1 × BIL6676) contains both *pui10.1* and *6.2*; IL12-3 contains above-mentioned *pui12.1*; F_1_ (IL10-1 × IL12-3) contains both *pui10.1* and *12.1*; BIL6546 contains both *pui6.2* and *12.1*; F_1_ (IL10-1 × BIL6546) contains all three *pui* loci *pui10.1*, *6.2* and *12.1*. Length of pollen tube growth is represented by circles (each circle corresponds a pollinated style as a biological replicate) in the plot, + indicates mean value of data. Different letters indicated significant differences (one-way ANOVA (Scheffe test), *P* < 0.05). Green, red and blue rectangles represent introgressed *pui6.2*,*10.1* and *12.1*, respectively. P/P and P/L in superscript indicate homozygous and heterozygous *pui* locus, respectively.

Compared with four genotypes of pollen tubes (M82, IL10-1, BIL6676 and F_1_ (IL10-1 × BIL6676)), pollen tubes from either F_1_ (IL10-1 × BIL6546) or BIL6546 grew significantly longer in LA0407 styles (Fig. 5a). In LA0407 styles, although a few pollen tubes of F_1_ (IL10-1 × IL12-3) and IL12-3 arrived at the bottom of LA0407 styles 72 HAP, no significant difference was found in pollen tube growth among F_1_ (IL10-1 × IL12-3), IL12-3, F_1_ (IL10-1 × BIL6676) and BIL6676 (Fig. 5a). In styles of LA1028, compared with pollen tubes of M82 and IL10-1, pollen tubes with introgressed *pui12.1*, such as IL12-3, BIL6546, F_1_ (IL10-1 × IL12-3) and F_1_ (IL10-1 × BIL6546), grew significantly longer 72 HAP (Fig. 5b). Taken together, these results suggest that *pui12.1* confers the ability of M82 pollen tubes to overcome the stylar response of LA0407 and LA1028, while further analysis of *pui10.1* and *6.2* are needed.

## Discussion

As an important interspecific RB, UI has received considerable attention in different genera (Covey *et al*., 2010, Takada *et al*., 2017, Yermishin *et al*., 2017, Lu *et al*., 2019). In *Solanum*, two major types of UI based on mating systems were observed: UI between 1) SI and SC species, and 2) two SC species (Pandey, 1962, Baek *et al*., 2015, Baek *et al*., 2016). In the analysis of UI between SC tomatoes, several sui factors have been identified, e.g. ODC2 (Qin & Chetelat, 2021), HT (Tovar-Méndez *et al*., 2017) and DIR1L (Muñoz-Sanz *et al*., 2021), but only one pui factor, FPS2, has been identified. It is thought that other pui factor(s), in addition to FPS2, present in the SC *S. pennellii* LA0716 (Qin *et al*., 2018). In this study, a *pui* locus at Chr. 6 (*pui6.2*) has been identified by forward genetics. During the functional validation of *pui6.2*, we found another *pui* locus (*pui12.1*) tightly linked to the *sui* locus *sui12.1*, which encodes HT proteins (Hamlin *et al*., 2017). Candidate genes were identified underlying *pui6.2* and *pui12.1*. Pollination assays demonstrated that three *pui* loci, *pui10.1*, *6.2* and *12.1*, functioned synergistically to break the stylar UI response of LA0716. Moreover, *pui12.1* is involved in breaking the stylar UI responses of *S. habrochaites* LA0407 and *S. chmielewskii* LA1028, suggesting an intersection of UI mechanisms in various tomato species.

In this study, ILs (Eshed & Zamir, 1995) and BILs (Ofner *et al*., 2016) between LA0716 and M82 were used for functional validation of *pui* loci. Interestingly, M82 pollen tubes containing homozygous *pui6.2*, *10.1* and *12.1* (F_2_ (BIL6546 × IL3-3)) grew to ∼87 % of the LA0716 stylar lengths *versus* M82 pollen tubes with attained only ∼27 % of the stylar length (Fig. 4), indicating that three *pui* loci are collectively required to break the stylar UI response in LA0716 styles. In addition to *pui6.2* and *12.1*, numerous genes are introgressed in the corresponding ILs or BILs as well. Therefore, further functional validation (e.g., transformation assay) of candidate genes in *pui6.2* and *12.1* is needed. However, once a *pui* factor is knocked out in the T_0_ generation of LA0716, it is difficult to obtain T_1_ generation with homozygous *pui* factor^loss-of-function^ (due to their self-sterility). Strategies, e.g., early bud pollination, can be used to overcome this challenge.

Previous research showed that HT proteins can be detected in pollen tubes (Goldraij *et al*., 2006), suggesting that there might be protein(s) in pollen tubes, which interact directly with HT either to exert its cytotoxicity for incompatibility or to detoxify HT in compatible interactions. Cysteine-rich peptides (CRPs) are characterized by a conserved N-terminal region containing a secretion peptide signal and a C-terminal cysteine-rich domain usually containing 4–16 cysteine residues (Marshall *et al*., 2011). Normally, two conserved cysteine motifs, i.e. CXXCXC and CXXXCC, are found at the C-terminus of the secreted HT-protein (Kondo *et al*., 2002a, Covey *et al*., 2010) (Fig. S4b), suggesting that HT proteins belong in the CRP category. Direct interactions between CRPs and its interacting proteins (especially receptor-like kinases (RLKs)) are reported in different pollen-pistil communications, e.g. the LAT52-LePRK2 complex (pollen tube hydration and germination) (Tang *et al*., 2002), SRK-SCR/SP11 (SI in stigma) (Murase *et al*., 2020), PrsS-PrpS (SI in style) (Wheeler *et al*., 2009), LURE-(MDIS1–MIKs) (pollen tube attraction) (Wang *et al*., 2016), RALFs-BUPSs (pollen tube rupture and sperm release) (Ge *et al*., 2017), PCP-Bs-FER (pollen tube hydration and germination) (Liu *et al*., 2021). However, in the Solanaceae, CRP-interacting proteins involved in pollen-style interactions remain undescribed. In this study, three HT-interacting proteins identified at *pui6.2* and *12.1* (Fig. S5 and S6) may have a potential function in detoxification against HT proteins.

One *pui* candidate gene, *Sopen06g027210*, encoding an ABC (ATP-binding cassette) transporter A family member (ABCA), is an ortholog of *Solyc06g070940* (*SlABCA5*) in *S. lycopersicum* (Fig. S7), which is specifically transcribed in mature tomato flowers (Ofori *et al*., 2018). ABC transporters in plants are grouped into seven subfamilies, namely ABCA to ABCG, and are responsible for the transmembrane transport of various compounds, e.g. coating materials, supportive materials, secondary metabolites and phytohormones (Hwang *et al*., 2016). A study in apple (*Malus domestica*) reported that an MdABCF transporter plays a role in SI by non-selectively transporting S-RNase into pollen tubes (Meng *et al*., 2014). In addition, ABC transporters are reported to sequester xenobiotics into vacuoles, such as during the detoxification of heavy mental ions (like cadmium and mercury) (Song *et al*., 2014). It is possible that the ABCA transporter encoded by *Sopen06g027210* plays a role in sequestration of HT proteins from pollen tubes. Further transgenic functional validation is needed to determine the function of *Sopen06g027210* in the pollen UI response.

Another candidate gene *Sopen06g026310* underlying *pui6.2* encodes an alpha subunit of the translation elongation factor (eEF1A), which is crucial for peptide-chain elongation during translation in eukaryotes (Andersen *et al*., 2003). The noncanonical functions of eEF1A were reported to promote replication or translation of certain RNA virus (e.g., tobacco mosaic virus) in plant cells (Chen *et al*., 2021). Further analysis is required to validate its role in UI responses.

At *pui12.1, Sopen12g029440* encoding a WD-repeat (WDR) domain containing protein is identified as another *pui* candidate gene. WDR proteins are characterized by several repeating units of ∼40-60 amino acids with C-terminal tryptophan (W)-aspartic acid (D) (namely WD40 domain). WD40 domains typically assemble into a β-propellers to provide binding sites for other proteins and foster transient interactions for these proteins (Jain & Pandey, 2018). In plants, WDR proteins are reported to be involved in a variety of cellular and developmental process, e.g., cell fate determination, cell division and cytokinesis, light signaling, flowering, and meristem organization (Airoldi *et al*., 2019, Schenk *et al*., 2021, Li *et al*., 2023a). Although the two homologous WDR proteins are supposed to play a role in UI, further validation (e.g., functional analysis of LA0716 pollen with double-knockout *pui* WDR genes) is required to elucidate their function.

In this study, two *pui* QTLs have been uncovered and functionally validated. Although further mechanistic studies are required for these *pui* factors, we are also interested in the evolutionary forces underlying the origin of these QTLs and related genes. Previous studies have demonstrated that SI factors, on both stylar and pollen sides of the interaction, are involved in UI between SI and SC species (Covey *et al*., 2010, Li & Chetelat, 2010, Tovar-Méndez *et al*., 2014, Li & Chetelat, 2015), suggesting that ecological or evolutionary forces that led to shifts in mating system may establish interspecific UI (Broz *et al*., 2017, Jewell *et al*., 2020). Two hypotheses to explain why genes contributing to UI become fixed in various tomato species are: 1) in wild tomatoes, genes involved in UI might be favored by natural selection and exhibit pleiotropic effect in the pollen-style interaction and 2) interspecific genetic conflict leading to UI might be due to genetic drift. Information about the ecological and molecular function of UI genes can inform us about the evolutionary forces underlying reproductive isolation and speciation.

## Acknowledgments

We thank Dr. Ben Zhang for providing plasmids and guidance for the mbSUS-based Y2H assays. We thank Dr. Shuqing Xu and Dr. Yuxing Xu for helpful discussions on bioinformatics analyses and manuscript drafting. We are grateful to the instrument and greenhouse support from the Service Center for Experimental Biotechnology at the Kunming Institute of Botany, CAS. This study was supported by the National Natural Science Foundation of China (32070346), Introducing Talents Fund of Chinese Academy of Sciences, Introducing Talents Start-up Fund of Kunming Institute of Botany, Chinese Academy of Sciences, Yunnan Revitalization Talent Support Program Young Talent Project (YNQR—QNRC—2020—118).

## Author contributions

WM, YL and HG designed research; HG contributed reagents and analytic tools; WM performed experimental work; YL performed bioinformatics work; WM and HG wrote the draft; YL, HM and ITB edited the draft. WM and YL contributed equally to this research.

## Data availability

The authors confirm that the data supporting the findings of this study are available within the article and its supplementary materials.

## Supporting information

**Table S1** Pollination results summarized from (Liedl *et al*., 1996, Hamlin *et al*., 2017, Qin *et al*., 2018, Qin & Chetelat, 2021).

**Table S2** Tomato materials used in the study.

**Table S3** DNA primers used in this study.

**Table S4** 55 differentially expressed genes (DEGs) between LA0716 and Heinz pollen linked to *pui10.1*.

**Table S5** 61 DEGs between LA0716 and Heinz pollen linked to *pui6.2*.

**Table S6** 13 candidate genes linked to *pui6.2*, which show at least a 2-fold difference in transcript abundance between LA0716 and Heinz pollen.

**Table S7** Introgression lines (Eshed & Zamir, 1995) or backcross inbred lines (Ofner *et al*., 2016) mentioned in this study and their introgression regions identified by SNP markers (Sim *et al*., 2012) or DNA resequencing.

**Table S8** *P*-values associated with patterns of transmission ratio distortion in 194 F_2_ plants of BIL6546 × IL3-3.

**Table S9** 12 candidate genes linked to *pui12.1*, which show at least a 2-fold difference in transcript abundance between LA0716 and Heinz pollen.

**Table S10** Genotype analysis to *Solyc12g056760* at *pui12.1* and its homologous gene *Solyc04g072120* in BC_1_ (Heinz × LA0716).

**Fig. S1** The Heinz genotype was not been detected in 155 progeny of LA0716 derived by binary mixture pollination.

**Fig. S2** UI response remains intact in introgression line 10-1 (IL10-1) harboring pollen UI factor *SpFPS2* in LA0716 styles.

**Fig. S3** Chromosome schematic diagram of IL6-2 and BIL6676.

**Fig. S4** Functional comparison of stylar UI factors between M82 and corresponding ILs/BILs.

**Fig. S5** Among candidate genes linked to *pui6.2*, significant differences in *Sopen06g027210* and *Sopen06g026310* transcript abundances between F_1_ (BIL6546 × IL3-3) and M82 pollen, and Y2H interactions with SpHT-A were found.

**Fig. S6** Among candidate genes linked to *pui12.1*, significant difference in *Sopen12g029440* transcript abundance was found in pollen between F_1_ (BIL6546 × IL3-3) and M82 pollen, and both its protein product and homologous protein encoded by *Sopen04g028290* directly interacted with SpHT-A in yeast.

**Fig. S7** Phylogenetic tree of ABC-A subfamily proteins of *S. pennellii*, *S. lycopersicum* and *Arabidopsis thaliana*.

