## Supplementary material for "Three pollen QTLs are sufficient to partially break stylar unilateral incompatibility of *Solanum pennellii*": Table S1, Table S2, Table S3, Table S4, Table S5, Table S6, Table S7, Table S8, Table S9, Table S10, Fig. S1, Fig. S2, Fig. S3, Fig. S4, Fig. S5, S6,

**This PDF file includes:**

Methods corresponding to the supplemental data

Table S1 to S10

Figure S1 to S7

References

### Methods corresponding to the supplemental data

#### Aniline blue staining to observe pollen tube growth

The growth of pollen tubes in styles was observed by aniline blue staining (Mori *et al.*, 2006). Images of a style or pistil were stitched by AutoStitch software (AutoStitch (matthewalunbrown.com). The length of the longest pollen tube in styles or pistils was measured by using ImageJ software (Jewell *et al.*, 2020, Li *et al.*, 2023). Phenotypes of compatible pollinations were repeated by more than 3 biological replicates. For incompatible pollinations, phenotypes were repeated by more than 6 biological replicates.

#### RNA-isolation, RNA-Seq and transcript profiling

Total RNA of pollen from Heinz and LA0716 was isolated using TRIzol™ Plus RNA Purification Kit (ThermoFisher Scientific, Cat. No. 12183555CN). 10 mg pollen of each sample was used for RNA extraction with 3 biological replicates. Library preparation and sequencing were conducted at the Annoroad company (Beijing, China). About 5 Gbp of clean data was generated from each replicate. RNA-Seq analysis was performed following the reported pipeline (Pertea *et al.*, 2016). Hisat2 (http://ccb.jhu.edu/software/hisat2) was used to align the reads to the tomato reference genome ITAG4.1 (https://solgenomics.net/organism/Solanum_lycopersicum/ genome). Transcript assembly and quantification were conducted using StringTie (http://ccb.jhu.edu/software/stringtie), where reference-guided mode was used to estimate transcript profiling. Ballgown (https://www.r-project.org) was used to identify differentially expressed genes (DEGs, $P < 0.05$ and $|\log_2\text{Fold-Change}| >= 1$).

#### Quantitative RT-PCR

RT-qPCR was carried out on the BIORAD CFX96 using the iTaq SYBR Green Universal mix (BIORAD, Cat. No. 1725125). The primer pairs used by RT-qPCR were listed in Table S3. The cDNA sequences of *eukaryotic translation initiation factor 5A-3* are identical in *S. lycopersicum* (*Solyc07g005560*) and *S. pennellii* (*Sopen07g001660*), which were used as the internal control.

**Mating-based split-ubiquitin (mbSUS) yeast two hybrid (Y2H) assay**

To detect interactions between SpHT-A and protein products of candidate genes at *pui6.2* and *12.1*, mbSUS-based Y2H assay was performed. The CDS sequence of SpHT-A without signal peptide (https://services.healthtech.dtu.dk/services/ SignalP-5.0/) was cloned into the *pExg2Met-Dest* vector to express the membrane-bound "bait" fusion protein with a glycosyl-phosphatidylinositol (GPI) signal sequence fused at the N-terminus (Zhang *et al.*, 2018). The candidate genes were cloned into a *pNX35-Dest* vector to express "prey" fusion proteins (Horaruang & Zhang, 2017). Primer pairs for cloning these candidate genes in mbSUS assay were listed in (Table S3). "Bait" and "prey" vectors were respectively transformed into THY. AP4 and THY. AP5 yeast strains and conducted the interaction verification as described previously (Horaruang & Zhang, 2017). Interaction between the ubiquitin N-terminal half NubG (mutated) with the C-terminal half Cub was represented as a negative control.

**Identification of ABC transporter A subfamily (ABCA) proteins in *S. pennellii* and phylogenetic tree construction**

ABC transporter A subfamily (ABCA) proteins in *S. pennellii* were identified as described previously (Ofori *et al.*, 2018). A genome-wide search for ABCA proteins in *S. pennellii* (https://solgenomics.net/organism/Solanum_pennellii/genome) was performed using the BLAST tool of Sol Genomics Network (SGN, http://www.solgenomics.net/). Based on ATP binding cassette (ABC) proteins reported previously in *S. lycopersicum* (ITAG3.10) (Ofori *et al.*, 2018), protein sequences of 8 SlABCA members were used as queries for BLAST search. Because *SlABCA8* (*Solyc03g113080*, ITAG3.10) (Ofori *et al.*, 2018) is not found in the genome of ITAG4.0 version (https://solgenomics.net/organism/Solanum_lycopersicum/ genome), it was not selected for BLAST search. Conserved domains of ABC transporter signature of predicted proteins were confirmed using InterPro web server (https://www.ebi.ac.uk/interpro/) (Paysan-Lafosse et al., 2023). Amino acid sequences of 8 SlABCA (*S. lycopersicum*), 8 SpABCA (*S. pennellii*) and 12 AtABCA (*Arabidopsis thaliana*, https://www.arabidopsis.org/) were used for phylogenetic analysis by MEGA11 software (https://www.megasoftware.net/). The sequences were aligned using the ClustalW algorithm. Following illustration previously reported (Tamura *et al.*, 2021), the phylogeny of these proteins was reconstructed using the maximum likelihood (ML) method by selecting the best-fit substitution model (JTT with Freqs. (+F) model) with 1000 bootstrap replicates.

**Supplemental tables**

**Table S1.** Pollination results summarized from (Qin *et al.*, 2018, Qin & Chetelat, 2021, Hamlin *et al.*, 2017, Liedl *et al.*, 1996).

| ♀ / ♂ | LA0716 | *S. lyc.* | $F_1$ (*S. lyc.* × LA0716) | IL3-3 | IL3-3$^{ODC2\text{-KD}}$ | LA0716$^{ODC2\text{-KD}}$ |
|---|---|---|---|---|---|---|
| LA0716 | + | + | + | + | + | + |
| LA0716$^{fps2}$ | - | + | - | - | + | + |
| *S. lycopersicum* (*S. lyc.*) | - | + | - | + | + | N.A. |
| $F_1$ (*S. lyc.* × LA0716$^{fps2}$) | N.A. | + | + | N.A. | N.A. | N.A. |

+: compatible pollination; -: incompatible pollination; LA0716$^{fps2}$: LA0716 with loss-of-function *SpFPS2*; IL3-3$^{ODC2\text{-KD}}$: IL3-3 with introgressed *SpODC2* genes that are knocked-down; LA0716$^{ODC2\text{-KD}}$: LA0716 with *SpODC2* genes that are knocked-down.

**Table S2.** Tomato materials used in the study.

| Tomato materials | | Accession number of TGRC |
|---|---|---|
| *S. pennellii* | | LA0716 |
| *S. habrochaites* | | LA0407 |
| *S. chmielewskii* | | LA1028 |
| *S. lycopersicum* | cv. Heinz | LA4345 |
| | cv. M82 | LA3475 |
| *S. pennellii* ILs/BILs (M82 genetic background) | IL3-3 | LA3488 |
| | IL10-1 | LA4087 |
| | IL12-2 | LA4099 |
| | IL12-3 | LA4100 |
| | IL12-4 | LA4102 |
| | BIL6676 | LA5213 |
| | BIL6546 | LA5113 |

**Table S3.** DNA primers used in this study.

| Gene ID | Primer name | Sequence (5' to 3') | Purpose |
|---|---|---|---|
| *Solyc07g005560* | eIF-5α-qF | GCAAGGTTGTTGAGGTCTCCACTT | RT-qPCR |
| *Sopen07g001660* | eIF-5α-qR | AGCTGATAGTCGGTACGGTTGACA | RT-qPCR |
| *Solyc03g098300* | ODC2-qF | CCGTTCGATTGCAGCAGAAA | RT-qPCR |
| *Sopen03g029020*<br>*Sopen03g029030*<br>*Sopen03g029040*<br>*Sopen03g029050* | ODC2-qR | TCAGTGTTGCATTTGACAGCA | RT-qPCR |
| *Solyc12g056585* | HT-A-qF | TTCACTTCCATTGCTTGAGGCA | RT-qPCR |
| *Sopen12g029190* | HT-A-qR | CATTTTCCACCTTCACCACCACCA | RT-qPCR |
| *Sopen12g029200* | HT-B-qF | GGATCATTGTTTCCTAATATAGCGTGT | RT-qPCR |
| *Sopen12g029200* | HT-B-qR | GTTTGAATTTGACTCAGATCACTCGT | RT-qPCR |
| *Solyc10g005820* | FPS2-qF | TCAGACTGCCTCTGGACAAA | RT-qPCR |
| *Sopen10g001780* | FPS2-qR | CACATGCCACTGGGAGATAA | RT-qPCR |
| *Solyc06g069520* | Sopen06g026810_qF | GCAAAGCCGTACAAGCCTTCG | RT-qPCR |
| *Sopen06g026810* | Sopen06g026810_qR | CACCAAAGACGCCGCAGATGA | RT-qPCR |
| *Solyc06g069020* | Sopen06g026310_qF | GTCCTTGATGCAGTCGAAGTTCCA | RT-qPCR |
| *Sopen06g026310* | Sopen06g026310_qR | CCTCACGTATGCTTCCAGACTCAA | RT-qPCR |
| *Solyc06g068720* | Sopen06g026000_qF | ACCAAGGTATGGCTGATGTGTTCC | RT-qPCR |
| *Sopen06g026000* | Sopen06g026000_qR | TGCTTGCTGCTGGCACAACTT | RT-qPCR |
| *Solyc06g067940* | Sopen06g02522_qF | CCAGGAGATGAAGGCGAAGACG | RT-qPCR |
| *Sopen06g025220* | Sopen06g025220_qR | GACCTCCTGATGGACACGACGA | RT-qPCR |
| *Solyc06g071770* | Sopen06g028060_qF | CAGTTGAGCCGGTGGTTGATGG | RT-qPCR |
| *Sopen06g028060* | Sopen06g028060_qR | ACTTCTGGTGCTGCCTCGGATA | RT-qPCR |
| *Solyc06g069410* | Sopen06g026690_qF | AGGCAATGTGGCTTCCGGTAGT | RT-qPCR |
| *Sopen06g026690* | Sopen06g026690_qR | ATCCTGAGCATCCATCGCCAGC | RT-qPCR |
| *Solyc06g071170* | Sopen06g027480_qF | TGGATGGAGATGGAGGTGATGA | RT-qPCR |
| *Sopen06g027480* | Sopen06g027480_qR | TGAAGCGAGAGGAACCACATTA | RT-qPCR |
| *Solyc06g066400* | Sopen06g021590_qF | GCTGACAGAACGATTGGACTGCTT | RT-qPCR |

| | | | |
|---|---|---|---|
| *Sopen06g021590* | Sopen06g021590_qR | ATGGCAGAATGATGACCCAGAAGG | RT-qPCR |
| *Solyc06g070920* | Sopen06g027200-qF | GGGTGTCCAGCGGATTTCAAGG | RT-qPCR |
| *Sopen06g027200* | Sopen06g027200-qR | GCTGCTGCTATCTCGCCTGTCT | RT-qPCR |
| *Solyc06g070940* | Sopen06g027210-qF | TGGATCACTTGGGAAGGACTCC | RT-qPCR |
| *Sopen06g027210* | Sopen06g027210-qR | TGTCGCTGAAGCTGCTTTGGT | RT-qPCR |
| *Solyc06g071020* | Sopen06g027320-qF | CGGAGCCTTCTTCGTGGAATCTG | RT-qPCR |
| *Sopen06g027320* | Sopen06g027320-qR | TGTTAGCCTGCCTGCGTCTGA | RT-qPCR |
| *Solyc06g068370* | Sopen06g025630-qF | TACATTCTGGTGTCAGTTGAGGATATTG | RT-qPCR |
| *Sopen06g025630* | Sopen06g025630-qR | ACATATTCAAGCTTCCAGGTCTCAATC | RT-qPCR |
| *Solyc06g071950* | Sopen06g028230-qF | GCTTCTCCTCCATGCCAATGTTGA | RT-qPCR |
| *Sopen06g028230* | Sopen06g028230-qR | GCTGTGCCAATCTGACTCATCCTC | RT-qPCR |
| *Solyc12g089150* | Sopen12g031020-qF | GATGCGGCGAGAACTGTTGAGA | RT-qPCR |
| *Sopen12g031020* | Sopen12g031020-qR | ACTGTCCATCTCCGTGTCCAAC | RT-qPCR |
| *Solyc12g088660* | Sopen12g030560-qF | AGCTCCCATGAGTGTTGCCTTG | RT-qPCR |
| *Sopen12g030560* | Sopen12g030560-qR | CCCTTTCGGAGGTGTTGGCTTT | RT-qPCR |
| *Solyc12g088180* | Sopen12g030180-qF | TTACCACGAGCAGCTCTCAGTTG | RT-qPCR |
| *Sopen12g030180* | Sopen12g030180-qR | CCACAGCAGCGTTGACATCCTT | RT-qPCR |
| *Solyc12g088980* | Sopen12g030880-qF | GCTGATGGCGTTGTCTATACCG | RT-qPCR |
| *Sopen12g030880* | Sopen12g030880-qR | CGCAGTCACAGCACACTCCT | RT-qPCR |
| *Solyc12g088200* | Sopen12g030200-qF | TGGTGGTGGTAGGCTATGCTCT | RT-qPCR |
| *Sopen12g030200* | Sopen12g030200-qR | CGACTCCACTTGCTTCCTGACA | RT-qPCR |
| *Solyc12g089310* | Sopen12g031170-qF | TCACTTGGAGGAGGGACTGGTT | RT-qPCR |
| *Sopen12g031170* | Sopen12g031170-qR | GGTGAGCATCATTCGGTCTGGA | RT-qPCR |
| *Solyc12g088950* | Sopen12g030850-qF | GCACGACGACAACTCTCAACAC | RT-qPCR |
| *Sopen12g030850* | Sopen12g030850-qR | AAAGCAGGCGAACAGAACACG | RT-qPCR |
| *Solyc12g057110* | Sopen12g029760-qF | CCGACTTGGACTGGCTCTCAAC | RT-qPCR |
| *Sopen12g029760* | Sopen12g029760-qR | CCAGTGTGTCCAACTCGGCAAT | RT-qPCR |
| *Solyc12g088360* | Sopen12g030340-qF | GGTGTTACGGCGGTGCTGAATT | RT-qPCR |
| *Sopen12g030340* | Sopen12g030340-qR | CGAAGCAGAGCACAAGCGGAAT | RT-qPCR |

| | | | |
|---|---|---|---|
| *Solyc12g088680* | Sopen12g030580-qF | TCCCTAGCAAACCGCCCAAGT | RT-qPCR |
| *Sopen12g030580* | Sopen12g030580-qR | GCCTCCACCCACTGTCTTCATT | RT-qPCR |
| *Solyc12g056550* | Sopen12g029160_qF | ACGCTTGTGGCTGCAATTCCT | RT-qPCR |
| *Sopen12g029160* | Sopen12g029160_qR | ACGCTCCCTCTTTCTTGACTCC | RT-qPCR |
| *Solyc12g056760* | Sopen12g029440-qF | GCCGAAGCGACTCCACTGTTAT | RT-qPCR |
| *Sopen12g029440* | Sopen12g029440-qR | TCCCACCCACCAAGGTTTGC | RT-qPCR |
| *Solyc04g072120* | Sopen04g028290-qF | GCCGAGGTGTTCCTATCAGATTCA | RT-qPCR |
| *Sopen04g028290* | Sopen04g028290-qR | GGTTGGATCAACGAGTCCTTAGGT | RT-qPCR |
| *Sopen03g029040* | SpODC2c-ILJD-F | ACTGAAATTGAAAAGACTCATC | Genotyping |
| *Sopen03g029040* | SpODC2c-ILJD-R | AATCACGAGTATCACATGG | Genotyping |
| *Solyc03g098300* | SlODC2-ILJD-F | TTTGAACAAAACCTGTTATCTCC | Genotyping |
| *Solyc03g098300* | SlODC2-ILJD-R | ATTTTACTAAGTATCACATGGGGA | Genotyping |
| *Sopen12g029190* | SpHT-ILJD-F | TGAAACTGTATTTACAGACAGT | Genotyping |
| *Sopen12g029190* | SpHT-ILJD-R | ATCGTCTTAAGGCGATGAAG | Genotyping |
| *Solyc12g056585* | SlHT-ILJD-F | AGCAAATCGTCTCATATTATCTTTAATC | Genotyping |
| *Solyc12g056585* | SlHT-ILJD-R | ATAATGATCCCCGCCTTTTC | Genotyping |
| *Solyc06g070940* | pui6.2-IDL-F | TTGACTGGGATTAATCCTGTTAC | Genotyping |
| *Sopen06g027210* | pui6.2-IDL-R | ATAGAGCTTCGTATGGACTG | Genotyping |
| *Solyc10g005820* | pui10.1-IDL-F | TAAGTGCACACATCCAAATTG | Genotyping |
| *Sopen10g001780* | pui10.1-IDL-R | TTCCAAAGCCAGATCACTG | Genotyping |
| *Solyc06g008710* | CUL1-IDL-F | CAGGAACGTGAGGGTGAGA | Genotyping |
| *Sopen06g003490* | CUL1-IDL-R | TCCACAAAAGTAACCCCTTCA | Genotyping |
| *Sopen06g027200* | Gene1tmd-attB1-F | GGGGACAAGTTTGTACAAAAAAGCAGGCTT CATGGCTGACTCCGTGACA | Y2H |
| *Sopen06g027200* | Gene1tmd-attB2-R | GGGGACCACTTTGTACAAGAAAGCTGGGTC ATTGATCTGGTCCAAGTAAAAAGC | Y2H |
| *Sopen06g027200* | Gene1cd-attB1-F | GGGGACAAGTTTGTACAAAAAAGCAGGCTT CATGTCATCAGGGAAAAGTCCCT | Y2H |
| *Sopen06g027200* | Gene1cd-attB2-F | GGGGACCACTTTGTACAAGAAAGCTGGGTC | Y2H |

| | | | |
|---|---|---|---|
| | | TGCTTCATTAAAGGATTGAGC | |
| *Sopen06g027210* | Gene2-attB1-F | GGGGACAAGTTTGTACAAAAAAGCAGGCTT CATGGAATTGCAGGGGGGA | Y2H |
| *Sopen06g027210* | Gene2-attB2-R | GGGGACCACTTTGTACAAGAAAGCTGGGTC TGACAAGTGCATTTTGAGTTGTG | Y2H |
| *Sopen06g027320* | Gene3-attB1-F | GGGGACAAGTTTGTACAAAAAAGCAGGCTT CATGGTAACTCTCATAAATAGGTCGAG | Y2H |
| *Sopen06g027320* | Gene3-attB2-R | GGGGACCACTTTGTACAAGAAAGCTGGGTC ACAAGGCTTCCCCTCAATG | Y2H |
| *Sopen06g026310* | Gene4-attB1-F | GGGGACAAGTTTGTACAAAAAAGCAGGCTT CATGGATATTGAAGAGGACATCAAG | Y2H |
| *Sopen06g026310* | Gene4-attB2-R | GGGGACCACTTTGTACAAGAAAGCTGGGTC TGCACTATCAGCAACAGTAG | Y2H |
| *Sopen06g026000* | Gene5-attB1-F | GGGGACAAGTTTGTACAAAAAAGCAGGCTT CATGTCAGTGGCTGGAGAG | Y2H |
| *Sopen06g026000* | Gene5-attB2-R | GGGGACCACTTTGTACAAGAAAGCTGGGTC ATCAAGATCTAGACTCTTTTTCATTG | Y2H |
| *Sopen06g026690* | Gene6-attB1-F | GGGGACAAGTTTGTACAAAAAAGCAGGCTT CATGGAAACAAGATCAGAAAAATTTTCTG | Y2H |
| *Sopen06g026690* | Gene6-attB2-R | GGGGACCACTTTGTACAAGAAAGCTGGGTC TTTGTTGAAGGCTCTTTGAGAC | Y2H |
| *Sopen06g021590* | Gene7-attB1-F | GGGGACAAGTTTGTACAAAAAAGCAGGCTT CATGGAATTAGCTGACAGAACG | Y2H |
| *Sopen06g021590* | Gene7-attB2-R | GGGGACCACTTTGTACAAGAAAGCTGGGTC TGCCTTCTTCTTTTTGGACTTG | Y2H |
| *Sopen12g029190* | Gene8-attB1-F | GGGGACAAGTTTGTACAAAAAAGCAGGCTT CGAAATGGTTGAGGCAAATCAAG | Y2H |
| *Sopen12g029190* | Gene8-attB2-R | GGGGACCACTTTGTACAAGAAAGCTGGGTC ACAACACATGGCTTTACAAACA | Y2H |

| | | | |
|---|---|---|---|
| *Sopen12g030180* | Gene9-attB1-F | GGGGACAAGTTTGTACAAAAAAGCAGGCTT<br>CATGAGAGAATGCATTTCAATCCAC | Y2H |
| *Sopen12g030180* | Gene9-attB2-R | GGGGACCACTTTGTACAAGAAAGCTGGGTC<br>GTACTCATCTCCTTCATCGCCA | Y2H |
| *Sopen12g029440* | Gene10-attB1-F | GGGGACAAGTTTGTACAAAAAAGCAGGCTT<br>CATGGTGGAGATCACAGAAG | Y2H |
| *Sopen12g029440* | Gene10-attB2-R | GGGGACCACTTTGTACAAGAAAGCTGGGTC<br>TTTTCGACGACTTTCCATG | Y2H |
| *Sopen12g030880* | Gene11-attB1-F | GGGGACAAGTTTGTACAAAAAAGCAGGCTT<br>CATGAAGGAATTTTCAGCTTT | Y2H |
| *Sopen12g030880* | Gene11-attB2-R | GGGGACCACTTTGTACAAGAAAGCTGGGTC<br>TTGAACATCTCCATTGTG | Y2H |
| *Sopen12g030200* | Gene12-attB1-F | GGGGACAAGTTTGTACAAAAAAGCAGGCTT<br>CATGAGGATGGGGGTTGAAAATG | Y2H |
| *Sopen12g030200* | Gene12-attB2-R | GGGGACCACTTTGTACAAGAAAGCTGGGTC<br>TTTCTGCTTCATAAGGCTCAG | Y2H |
| *Sopen12g030850* | Gene13-attB1-F | GGGGACAAGTTTGTACAAAAAAGCAGGCTT<br>CATGCAGGTGTATCGATGTTTTC | Y2H |
| *Sopen12g030850* | Gene13-attB2-R | GGGGACCACTTTGTACAAGAAAGCTGGGTC<br>ATATCCTTGGAAGATTGGTGG | Y2H |

**Table S4.** 55 differentially expressed genes (DEGs) between LA0716 and Heinz pollen linked to *pui10.1*.

| Gene ID | | Physical location in Chr.10 (bp) | | Difference of paternal genotype ratio (P – L) | Fold change | *P*-value | Average TPM in pollen | |
|---|---|---|---|---|---|---|---|---|
| Heinz | LA0716 | Start | End | | | | Heinz | LA0716 |
| *Solyc10g005230* | *Sopen10g001220* | 61926 | 66510 | 0.93 | 0.25 | 0.0038 | 28.46 | 8.53 |
| *Solyc10g005560* | *Sopen10g001230* | 349603 | 353904 | 0.98 | 2.49 | 0.0285 | 9.26 | 24.85 |
| *Solyc10g005580* | *Sopen10g001240* | 355545 | 360396 | 0.98 | 0.35 | 0.0200 | 7.98 | 1.70 |
| *Solyc10g005590* | *Sopen10g001250* | 361236 | 368068 | 0.97 | 0.52 | 0.0414 | 2.98 | 1.36 |
| *Solyc10g005600* | *Sopen10g001260* | 368582 | 376779 | 0.97 | 4.97 | 0.0003 | 0.10 | 4.43 |
| *Solyc10g005620* | *Sopen10g001270* | 386956 | 390817 | 0.96 | 1.92 | 0.0339 | 0.25 | 1.77 |
| *Solyc10g005650* | *Sopen10g001280* | 405887 | 419153 | 0.98 | 3.23 | 0.0041 | 4.94 | 22.83 |
| *Solyc10g005670* | *Sopen10g001290* | 428605 | 428883 | 0.98 | 0.22 | 0.0002 | 60.99 | 10.74 |
| *Solyc10g005700* | *Sopen10g001670* | 453482 | 457532 | 0.98 | 0.54 | 0.0075 | 2.12 | 1.20 |
| *Solyc10g005720* | *Sopen10g001680* | 462713 | 463024 | 0.98 | 32.31 | 0.0010 | 0.14 | 34.05 |
| *Solyc10g005750* | *Sopen10g001720* | 493657 | 498019 | 0.96 | 0.36 | 0.0029 | 8.03 | 2.31 |
| *Solyc10g005800* | *Sopen10g001770* | 519303 | 524168 | 0.98 | 0.40 | 0.0052 | 5.74 | 3.68 |
| *Solyc10g005820_FPS2* | *Sopen10g001780* | 525499 | 531017 | 0.98 | 86.29 | 0.0000 | 0.72 | 131.31 |
| *Solyc10g005850* | *Sopen10g001810* | 543156 | 547995 | 0.99 | 0.38 | 0.0091 | 16.42 | 6.24 |
| *Solyc10g005860* | *Sopen10g001810* | 548472 | 550348 | 0.97 | 0.34 | 0.0043 | 8.24 | 3.36 |
| *Solyc10g006070* | *Sopen10g002020* | 685526 | 689081 | 0.98 | 1.39 | 0.0345 | 1.01 | 2.77 |
| *Solyc10g006090* | *Sopen10g002040* | 700591 | 711911 | 0.97 | 1.52 | 0.0015 | 2.15 | 4.07 |
| *Solyc10g006110* | *Sopen10g002060* | 733370 | 738658 | 0.96 | 0.32 | 0.0042 | 443.75 | 148.19 |
| *Solyc10g006130* | *Sopen10g002080* | 750250 | 750915 | 0.97 | 5.06 | 0.0004 | 61.37 | 337.19 |
| *Solyc10g006170* | *Sopen10g002120* | 770550 | 773792 | 0.98 | 0.34 | 0.0001 | 1.78 | 0.03 |
| *Solyc10g006210* | *Sopen10g002170* | 795987 | 796313 | 0.96 | 0.33 | 0.0219 | 2.21 | 0.13 |
| *Solyc10g006230* | *Sopen10g002190* | 806192 | 810040 | 0.95 | 0.39 | 0.0458 | 1.44 | 1.45 |
| *Solyc10g006240* | *Sopen10g002200* | 811803 | 816356 | 0.96 | 3.09 | 0.0063 | 6.07 | 22.46 |
| *Solyc10g006260* | *Sopen10g002210* | 818467 | 821686 | 0.97 | 0.56 | 0.0071 | 32.28 | 22.47 |
| *Solyc10g006480* | *Sopen10g002430* | 957680 | 960486 | 0.98 | 1.22 | 0.0177 | 31.46 | 42.51 |
| *Solyc10g006490* | *Sopen10g002440* | 961839 | 965953 | 0.99 | 2.08 | 0.0004 | 1.25 | 3.76 |
| *Solyc10g006500* | *Sopen10g002450* | 967777 | 972001 | 0.99 | 0.38 | 0.0073 | 1.61 | 0.02 |
| *Solyc10g006600* | *Sopen10g002540* | 1009604 | 1021646 | 0.98 | 0.16 | 0.0014 | 10.69 | 0.60 |
| *Solyc10g006610* | *Sopen10g002550* | 1020531 | 1023831 | 0.98 | 0.12 | 0.0001 | 10.08 | 0.41 |
| *Solyc10g006730* | *Sopen10g002650* | 1076355 | 1078088 | 0.98 | 0.08 | 0.0008 | 16.38 | 0.55 |
| *Solyc10g006770* | *Sopen10g002690* | 1104426 | 1107940 | 0.99 | 1.81 | 0.0091 | 1.60 | 3.68 |
| *Solyc10g006850* | *Sopen10g002760* | 1169455 | 1176565 | 0.96 | 2.41 | 0.0199 | 7.54 | 14.90 |
| *Solyc10g006930* | *Sopen10g002840* | 1245060 | 1256917 | 0.95 | 0.60 | 0.0255 | 1.74 | 1.55 |
| *Solyc10g006960* | *Sopen10g002870* | 1268908 | 1269609 | 0.96 | 3.50 | 0.0007 | 13.93 | 45.19 |
| *Solyc10g007010* | *Sopen10g002920* | 1297226 | 1297465 | 0.96 | 4.39 | 0.0086 | 1.36 | 10.64 |
| *Solyc10g007040* | *Sopen10g002940* | 1315652 | 1318419 | 0.95 | 0.81 | 0.0141 | 1.55 | 1.30 |
| *Solyc10g007140* | *Sopen10g003040* | 1427510 | 1431711 | 0.93 | 2.44 | 0.0218 | 54.86 | 129.10 |

| | | | | | | | | |
|---|---|---|---|---|---|---|---|---|
| *Solyc10g007156* | *Sopen10g003350* | 1478911 | 1479830 | 0.92 | 2.91 | 0.0185 | 0.00 | 1.44 |
| *Solyc10g007190* | *Sopen10g003290* | 1509100 | 1511802 | 0.91 | 0.39 | 0.0222 | 1.73 | 0.22 |
| *Solyc10g007200* | *Sopen10g003280* | 1512234 | 1517689 | 0.90 | 2.41 | 0.0003 | 0.07 | 1.64 |
| *Solyc10g007250* | *Sopen10g003070* | 1547195 | 1549140 | 0.95 | 0.35 | 0.0030 | 1.82 | 0.00 |
| *Solyc10g007290* | *Sopen10g003110* | 1566772 | 1574945 | 0.93 | 8.93 | 0.0003 | 3.15 | 35.95 |
| *Solyc10g007320* | *Sopen10g003130* | 1603822 | 1609680 | 0.94 | 0.54 | 0.0163 | 1.17 | 0.28 |
| *Solyc10g007410* | *Sopen10g003240* | 1653457 | 1660385 | 0.92 | 1.92 | 0.0245 | 10.26 | 22.51 |
| *Solyc10g007420* | *Sopen10g003250* | 1666582 | 1669511 | 0.94 | 4.31 | 0.0003 | 9.24 | 38.21 |
| *Solyc10g007530* | *Sopen10g003470* | 1731970 | 1733450 | 0.90 | 2.98 | 0.0127 | 15.09 | 37.47 |
| *Solyc10g007680* | *Sopen10g003590* | 1825716 | 1833212 | 0.92 | 0.57 | 0.0045 | 1.04 | 0.17 |
| *Solyc10g007730* | *Sopen10g003640* | 1857634 | 1860612 | 0.91 | 0.07 | 0.0009 | 28.36 | 1.42 |
| *Solyc10g007740* | *Sopen10g003650* | 1861695 | 1867891 | 0.92 | 0.45 | 0.0011 | 1.47 | 0.52 |
| *Solyc10g007780* | *Sopen10g003690* | 1895297 | 1896004 | 0.91 | 0.50 | 0.0471 | 7.03 | 2.48 |
| *Solyc10g007810* | *Sopen10g003710* | 1908241 | 1913674 | 0.90 | 1.65 | 0.0063 | 0.49 | 1.31 |
| *Solyc10g008110* | *Sopen10g003980* | 2117846 | 2124970 | 0.90 | 1.79 | 0.0015 | 2.78 | 8.19 |
| *Solyc10g008120* | *Sopen10g003990* | 2143926 | 2145899 | 0.88 | 15.60 | 0.0015 | 0.09 | 37.48 |
| *Solyc10g008300* | *Sopen10g004160* | 2323474 | 2328157 | 0.89 | 3.59 | 0.0200 | 7.61 | 29.19 |
| *Solyc10g008390* | *Sopen10g004220* | 2400999 | 2405504 | 0.88 | 2.28 | 0.0028 | 1.61 | 4.43 |

**Table S5.** 61 DEGs between LA0716 and Heinz pollen linked to *pui6.2*.

| Gene ID | | Physical location in Chr.6 (bp) | | Difference of paternal genotype ratio (P – L) | Fold change | *P*-value | Average TPM in pollen | |
|---|---|---|---|---|---|---|---|---|
| Heinz | LA0716 | Start | End | | | | Heinz | LA0716 |
| *Solyc06g066330* | *Sopen06g021690* | 39251327 | 39256766 | 0.87 | 0.58 | 0.0260 | 1.52 | 1.13 |
| *Solyc06g066400* | *Sopen06g021590* | 39319019 | 39321319 | 0.88 | 0.22 | 0.0016 | 24.99 | 5.73 |
| *Solyc06g066410* | *Sopen06g024700* | 39327050 | 39328343 | 0.86 | 31.89 | 0.0019 | 3.32 | 109.04 |
| *Solyc06g066440* | *Sopen06g024730* | 39344700 | 39349379 | 0.87 | 1.85 | 0.0177 | 24.17 | 39.22 |
| *Solyc06g066550* | *Sopen06g024820* | 39440266 | 39446891 | 0.86 | 2.69 | 0.0042 | 0.73 | 3.47 |
| *Solyc06g066700* | *Sopen06g024990* | 39546658 | 39546906 | 0.87 | 12.92 | 0.0002 | 0.00 | 10.77 |
| *Solyc06g067940* | *Sopen06g025220* | 39746987 | 39750282 | 0.86 | 5.43 | 0.0062 | 6.43 | 38.63 |
| *Solyc06g068230* | *Sopen06g025510* | 39920341 | 39930856 | 0.86 | 0.21 | 0.0072 | 3.80 | 1.15 |
| *Solyc06g068370* | *Sopen06g025630* | 39993996 | 39998452 | 0.87 | 0.19 | 0.0002 | 21.20 | 3.80 |
| *Solyc06g068460* | *Sopen06g025710* | 40068447 | 40070421 | 0.88 | 1.75 | 0.0436 | 0.45 | 2.92 |
| *Solyc06g068710* | *Sopen06g025990* | 40228975 | 40229873 | 0.88 | 2.87 | 0.0041 | 1.39 | 3.92 |
| *Solyc06g068720* | *Sopen06g026000* | 40231224 | 40238198 | 0.88 | 3.28 | 0.0215 | 5.11 | 19.36 |
| *Solyc06g068780* | *Sopen06g026060* | 40261477 | 40266085 | 0.88 | 0.18 | 0.0073 | 6.69 | 0.93 |
| *Solyc06g068900* | *Sopen06g026190* | 40351353 | 40353054 | 0.90 | 1.49 | 0.0148 | 48.23 | 72.07 |
| *Solyc06g068990* | *Sopen06g026280* | 40441977 | 40448095 | 0.89 | 2.54 | 0.0391 | 1.28 | 4.13 |
| *Solyc06g069020* | *Sopen06g026310* | 40456757 | 40468804 | 0.88 | 2.38 | 0.0177 | 4.14 | 12.80 |
| *Solyc06g069380* | *Sopen06g026640* | 40760250 | 40763960 | 0.89 | 0.62 | 0.0184 | 2.30 | 1.04 |
| *Solyc06g069400* | *Sopen06g026670* | 40792184 | 40795349 | 0.89 | 2.06 | 0.0016 | 0.00 | 1.01 |
| *Solyc06g069410* | *Sopen06g026690* | 40796453 | 40801404 | 0.90 | 0.33 | 0.0013 | 230.73 | 84.22 |
| *Solyc06g069450* | *Sopen06g026740* | 40832451 | 40836852 | 0.90 | 1.34 | 0.0417 | 0.56 | 1.29 |
| *Solyc06g069520* | *Sopen06g026810* | 40923182 | 40925930 | 0.92 | 2.58 | 0.0029 | 14.98 | 46.01 |
| *Solyc06g069560* | *Sopen06g026850* | 40942891 | 40944138 | 0.91 | 1.67 | 0.0364 | 25.17 | 45.00 |
| *Solyc06g070920* | *Sopen06g027200* | 41220142 | 41223563 | 0.93 | 2.31 | 0.0293 | 688.20 | 1688.29 |
| *Solyc06g070940* | *Sopen06g027210* | 41226357 | 41231370 | 0.93 | 5.99 | 0.0037 | 400.74 | 2533.21 |
| *Solyc06g070950* | *Sopen06g027220* | 41232015 | 41240070 | 0.92 | 1.72 | 0.0121 | 0.03 | 1.27 |
| *Solyc06g071020* | *Sopen06g027320* | 41289597 | 41291616 | 0.92 | 2.28 | 0.0218 | 203.65 | 385.02 |
| *Solyc06g071070* | *Sopen06g027370* | 41328092 | 41328877 | 0.91 | 0.42 | 0.0055 | 1.06 | 0.28 |
| *Solyc06g071110* | *Sopen06g027410* | 41358372 | 41365502 | 0.90 | 1.88 | 0.0338 | 9.53 | 16.60 |
| *Solyc06g071140* | *Sopen06g027440* | 41395256 | 41402346 | 0.90 | 0.42 | 0.0290 | 3.54 | 1.44 |
| *Solyc06g071170* | *Sopen06g027480* | 41423134 | 41433077 | 0.89 | 0.48 | 0.0420 | 26.29 | 14.13 |
| *Solyc06g071230* | *Sopen06g027540* | 41478463 | 41480884 | 0.90 | 1.57 | 0.0286 | 5.50 | 9.85 |
| *Solyc06g071250* | *Sopen06g027560* | 41497291 | 41506946 | 0.90 | 0.49 | 0.0107 | 1.27 | 0.08 |
| *Solyc06g071290* | *Sopen06g027590* | 41521314 | 41526488 | 0.90 | 0.09 | 0.0003 | 19.75 | 0.87 |
| *Solyc06g071470* | *Sopen06g027750* | 41635100 | 41641866 | 0.87 | 2.20 | 0.0454 | 1.16 | 5.39 |
| *Solyc06g071550* | *Sopen06g027840* | 41714191 | 41718072 | 0.87 | 0.47 | 0.0032 | 1.72 | 0.87 |
| *Solyc06g071770* | *Sopen06g028060* | 41866452 | 41870790 | 0.86 | 2.28 | 0.0081 | 11.05 | 25.15 |
| *Solyc06g071850* | *Sopen06g028130* | 41928446 | 41936591 | 0.86 | 0.50 | 0.0081 | 3.58 | 1.16 |
| *Solyc06g071870* | *Sopen06g028150* | 41944131 | 41946708 | 0.86 | 0.51 | 0.0352 | 1.92 | 1.04 |
| *Solyc06g071950* | *Sopen06g028230* | 41989995 | 41990948 | 0.86 | 0.15 | 0.0044 | 17.98 | 1.91 |

| | | | | | | | | |
|---|---|---|---|---|---|---|---|---|
| *Solyc06g072140* | *Sopen06g028440* | 42129504 | 42135893 | 0.86 | 0.63 | 0.0219 | 3.41 | 2.53 |
| *Solyc06g072510* | *Sopen06g028880* | 42382730 | 42384934 | 0.86 | 4.67 | 0.0396 | 35.30 | 130.42 |
| *Solyc06g072530* | *Sopen06g028900* | 42407363 | 42408478 | 0.83 | 8.92 | 0.0004 | 0.24 | 10.53 |
| *Solyc06g072590* | *Sopen06g028940* | 42434827 | 42436510 | 0.85 | 0.72 | 0.0208 | 533.35 | 375.36 |
| *Solyc06g072600* | *Sopen06g028950* | 42437247 | 42439175 | 0.86 | 0.30 | 0.0209 | 2.00 | 0.55 |
| *Solyc06g072740* | *Sopen06g029120* | 42527289 | 42528242 | 0.85 | 1.97 | 0.0010 | 23.87 | 34.52 |
| *Solyc06g072770* | *Sopen06g029150* | 42538247 | 42542189 | 0.85 | 1.68 | 0.0466 | 42.74 | 59.57 |
| *Solyc06g072790* | *Sopen06g029170* | 42551144 | 42559785 | 0.86 | 1.48 | 0.0351 | 1.02 | 2.15 |
| *Solyc06g072920* | *Sopen06g029280* | 42617609 | 42619107 | 0.85 | 2.05 | 0.0149 | 12.83 | 23.72 |
| *Solyc06g073020* | *Sopen06g029370* | 42661369 | 42663802 | 0.83 | 0.63 | 0.0138 | 3.52 | 2.16 |
| *Solyc06g073080* | *Sopen06g029500* | 42683039 | 42685997 | 0.81 | 0.53 | 0.0259 | 152.44 | 68.15 |
| *Solyc06g073150* | *Sopen06g029430* | 42729711 | 42734560 | 0.82 | 1.82 | 0.0480 | 2.76 | 5.50 |
| *Solyc06g073160* | *Sopen06g029510* | 42736160 | 42742007 | 0.82 | 2.19 | 0.0065 | 11.33 | 26.45 |
| *Solyc06g073190* | *Sopen06g029560* | 42758420 | 42761191 | 0.82 | 4.73 | 0.0063 | 18.73 | 94.84 |
| *Solyc06g073230* | *Sopen06g029590* | 42787788 | 42796092 | 0.82 | 0.49 | 0.0230 | 3.50 | 1.68 |
| *Solyc06g073300* | *Sopen06g029660* | 42825164 | 42825571 | 0.82 | 0.43 | 0.0412 | 5.65 | 4.00 |
| *Solyc06g073310* | *Sopen06g029670* | 42827684 | 42829886 | 0.81 | 0.34 | 0.0029 | 17.85 | 6.86 |
| *Solyc06g073330* | *Sopen06g029710* | 42846033 | 42853792 | 0.82 | 0.49 | 0.0042 | 2.14 | 1.19 |
| *Solyc06g073340* | *Sopen06g029720* | 42856980 | 42864743 | 0.82 | 0.61 | 0.0406 | 0.86 | 1.57 |
| *Solyc06g073470* | *Sopen06g029830* | 42945077 | 42950612 | 0.81 | 2.82 | 0.0020 | 0.15 | 2.24 |
| *Solyc06g073650* | *Sopen06g030060* | 43110545 | 43112167 | 0.80 | 0.64 | 0.0055 | 1.85 | 1.26 |
| *Solyc06g073700* | *Sopen06g030100* | 43133695 | 43136747 | 0.80 | 0.48 | 0.0080 | 3.65 | 3.06 |

**Table S6.** 13 candidate genes linked to *pui6.2*, which show at least a 2-fold difference in transcript abundance between LA0716 and Heinz pollen.

| Gene ID (Heinz) | Gene ID (LA0716) | Annotation | Difference of paternal genotype ratio (P – L) | Fold change | *P*-value | Average TPM in pollen | |
|---|---|---|---|---|---|---|---|
| | | | | | | Heinz | LA0716 |
| **Candidate genes with a higher transcript abundance in LA0716 pollen compared to Heinz pollen** | | | | | | | |
| *Solyc06g070920* | *Sopen06g027200* | ABC transporter A family member 7 | 0.93 | 2.45 | 0.0154 | 688.20 | 1688.29 |
| *Solyc06g070940* | *Sopen06g027210* | ABC transporter A family member 2 | 0.93 | 6.32 | 0.0024 | 400.74 | 2533.21 |
| *Solyc06g071020* | *Sopen06g027320* | Pectate lyase | 0.92 | 2.28 | 0.0218 | 203.65 | 385.02 |
| *Solyc06g069520* | *Sopen06g026810* | Protein Asterix | 0.92 | 3.07 | 0.0053 | 14.98 | 46.01 |
| *Solyc06g069020* | *Sopen06g026310* | Eukaryotic peptide chain release factor GTP-binding subunit ERF3A | 0.90 | 3.09 | 0.0025 | 4.14 | 12.80 |
| *Solyc06g068720* | *Sopen06g026000* | Mitochondrial substrate carrier family protein | 0.89 | 3.79 | 0.0007 | 5.11 | 19.36 |
| *Solyc06g067940* | *Sopen06g025220* | Cytochrome c oxidase subunit 5b-1 | 0.86 | 5.43 | 0.0062 | 6.43 | 38.63 |
| *Solyc06g071770* | *Sopen06g028060* | PB1 domain protein | 0.86 | 2.28 | 0.0081 | 11.05 | 25.15 |
| **Candidate genes with a lower transcript abundance in LA0716 pollen compared to Heinz pollen** | | | | | | | |
| *Solyc06g069410* | *Sopen06g026690* | ADP/ATP carrier protein | 0.90 | 0.33 | 0.0013 | 230.73 | 84.22 |
| *Solyc06g071170* | *Sopen06g027480* | tRNA-lysidine synthase | 0.89 | 0.48 | 0.0420 | 26.29 | 14.13 |
| *Solyc06g066400* | *Sopen06g021590* | Dolichol phosphate-mannose biosynthesis regulatory protein | 0.88 | 0.22 | 0.0016 | 24.99 | 5.73 |
| *Solyc06g068370* | *Sopen06g025630* | Unknown protein | 0.87 | 0.19 | 0.0002 | 21.20 | 3.80 |
| *Solyc06g071950* | *Sopen06g028230* | GDP-mannose transporter GONST3 | 0.86 | 0.15 | 0.0044 | 17.98 | 1.91 |

**Table S7**. Introgression lines (Eshed & Zamir, 1995) or backcross inbred lines (Ofner *et al.*, 2016) mentioned in this study and their introgression regions identified by SNP markers (Sim *et al.*, 2012) or DNA resequencing.

| SNP marker | Mb | Chromosome | Introgression lines or backcross inbred lines introgressed with segments from LA0716 | | | |
|---|---|---|---|---|---|---|
| *solcap_snp_sl_45992* | 0.05 | Chr.10 | IL10-1 | | | |
| *SL10670_292* | 79.3 | Chr.10 | IL10-1 | | | |
| *solcap_snp_sl_8774* | 122.8 | Chr.10 | | | | |
| *CL017080-0343* | 0.31 | Chr.3 | | | | |
| *solcap_snp_sl_32530* | 47.5 | Chr.3 | IL3-3 | | | |
| *SGN-U579212_snp25451* | 84.9 | Chr.3 | IL3-3 | | | |
| *solcap_snp_sl_67626* | 121.7 | Chr.3 | | | | |
| *solcap_snp_sl_39981* | 0.06 | Chr.6 | | | BIL6546 | |
| *CL017670-0525* | 35.73 | Chr.6 | IL6-2 | | BIL6546 | |
| *CL009178-0553* | 37.32 | Chr.6 | IL6-2 | | | |
| *solcap_snp_sl_56058* | 39.28 | Chr.6 | IL6-2 | BIL6676 | | |
| *solcap_snp_sl_57766* | 40.39 | Chr.6 | IL6-2 | BIL6676 | **BIL6546**$^{+}$ | |
| *solcap_snp_sl_57327* | 42.5 | Chr.6 | IL6-2 | BIL6676 | **BIL6546**$^{+}$ | |
| *CL016102-0429_solcap_snp_sl_57352* | 45.95 | Chr.6 | IL6-2 | BIL6676 | | |
| *solcap_snp_sl_54183* | 48.8 | Chr.6 | | | | |
| *solcap_snp_sl_28840* | 0.05 | Chr.12 | | | | |
| *solcap_snp_sl_17703* | 0.09 | Chr.12 | | IL12-2 | | |
| *solcap_snp_sl_41031* | 2.33 | Chr.12 | BIL6546 | IL12-2 | | |
| *solcap_snp_sl_41162* | 3.03 | Chr.12 | BIL6546 | IL12-2 | IL12-3 | |
| *solcap_snp_sl_19344* | 61.86 | Chr.12 | BIL6546 | IL12-2 | IL12-3 | |
| *solcap_snp_sl_14423* | 63.68 | Chr.12 | BIL6546 | | IL12-3 | |
| *solcap_snp_sl_31966* | 64.06 | Chr.12 | BIL6546 | | IL12-3 | IL12-4 |
| *solcap_snp_sl_25007* | 64.81 | Chr.12 | BIL6546 | | IL12-3 | IL12-4 |
| *solcap_snp_sl_31585* | 66 | Chr.12 | BIL6546 | | IL12-3 | IL12-4 |
| *solcap_snp_sl_31342* | 66.89 | Chr.12 | | | | IL12-4 |

**BIL6546**$^{+}$: the introgressed region of 40.39–42.5 Mbp on Chr. 6 was validated by DNA resequencing.

**Table S8.** *P*-values associated with patterns of transmission ratio distortion in 194 $F_2$ plants of BIL6546 × IL3-3.

| Chr.3_*ODC2* | Chr.12_*HT*-A | | | Chr.6_*pui6.2* |
|---|---|---|---|---|
| | LL | PL | PP | |
| LL | 0.04711 | 0.00211 | 0.41442 | LL |
| | 0.04118 | 0.11799 | 0.01125 | PL |
| | 0.22223 | 0.01125 | 0.022 | PP |
| PL | 0.00211 | 0.0003 | 0.04118 | LL |
| | 0.00413 | 0.0796 | 0.0027 | PL |
| | 0.08502 | 0.09492 | 1.8E-07 | PP |
| PP | 0.04711 | 0.01323 | 0.14508 | LL |
| | 0.00211 | 0.00934 | 0.10655 | PL |
| | 0.14508 | 0.04239 | 0.00005 | PP |

LL, homozygous *S. lycopersicum* genotype; LP, heterozygous genotype; PP, homozygous *S. pennellii* genotype.

**Table S9.** 12 candidate genes linked to *pui12.1*, which show at least a 2-fold difference in transcript abundance between LA0716 and Heinz pollen.

| Gene ID (Heinz) | Gene ID (LA0716) | Annotation | Difference of paternal genotype ratio (P – L) | Fold change | *P*-value | Average TPM in pollen | |
|---|---|---|---|---|---|---|---|
| | | | | | | Heinz | LA0716 |
| **Candidate genes with a higher transcript abundance in LA0716 pollen compared to Heinz pollen** | | | | | | | |
| *Solyc12g088200* | *Sopen12g030200* | Inositol-tetrakisphosphate 1-kinase | 0.78 | 2.13 | 0.0495 | 125.60 | 280.05 |
| *Solyc12g089310* | *Sopen12g031170* | Tubulin beta chain | 0.77 | 2.13 | 0.0125 | 41.33 | 70.12 |
| *Solyc12g088180* | *Sopen12g030180* | Tubulin alpha chain | 0.76 | 474.76 | 0.0000 | 0.05 | 440.34 |
| *Solyc12g088980* | *Sopen12g030880* | Transducin/WD40 repeat-like protein | 0.76 | 4.84 | 0.0021 | 76.81 | 387.49 |
| *Solyc12g056760* | *Sopen12g029440* | WD repeat protein | 0.76 | 3.58 | 0.0080 | 84.41 | 315.27 |
| *Solyc12g088950* | *Sopen12g030850* | Leucine-rich repeat family protein | 0.76 | 2.32 | 0.0071 | 83.23 | 177.97 |
| *Solyc12g057110* | *Sopen12g029760* | 14-3-3 protein | 0.74 | 2.24 | 0.0008 | 707.83 | 1701.22 |
| **Candidate genes with a lower transcript abundance in LA0716 pollen compared to Heinz pollen** | | | | | | | |
| *Solyc12g088360* | *Sopen12g030340* | RING-type E3 ubiquitin transferase | 0.78 | 0.36 | 0.0014 | 36.81 | 10.48 |
| *Solyc12g088680* | *Sopen12g030580* | Ubiquitin-conjugating enzyme | 0.77 | 0.24 | 0.0084 | 18.99 | 4.14 |
| *Solyc12g056550* | *Sopen12g029160* | Plant/F1M20-13 protein | 0.77 | 0.43 | 0.0075 | 76.74 | 35.86 |
| *Solyc12g089150* | *Sopen12g031020* | Syntaxin-61 | 0.76 | 0.27 | 0.0015 | 24.99 | 6.00 |
| *Solyc12g088660* | *Sopen12g030560* | Mannose-P-dolichol utilization defect protein | 0.76 | 0.21 | 0.0031 | 10.36 | 2.16 |

**Table S10.** Genotype analysis to *Solyc12g056760* at *pui12.1* and its homologous gene *Solyc04g072120* in $BC_1$ (Heinz × LA0716).

| *Solyc04g072120/Sopen04g028290* (WDR_Chr. 04) | *Solyc12g056760/Sopen12g029440* (WDR_Chr. 12) | | Total |
|---|---|---|---|
| | PP | PL | |
| PP | 45 (25) | 15 (25) | 60 (50) |
| PL | 32 (25) | 8 (25) | 40 (50) |
| Total | 77 (50) | 23 (50) | 100 (100) |
| $X^2_{WDR\_Chr.\ 04}$ | 4 | 0.0412 | |
| $X^2_{WDR\_Chr.\ 12}$ | 29.16 | 1.40E-10 | |
| $X^2_{WDR\_Chr.\ 04\ \&\ 12}$ | 33.52 | 1.65231E-11 | |

LL, homozygous *S. lycopersicum* genotype; LP, heterozygous genotype; PP, homozygous *S. pennellii* genotype. Expected values are in parentheses.

### Supplemental figures

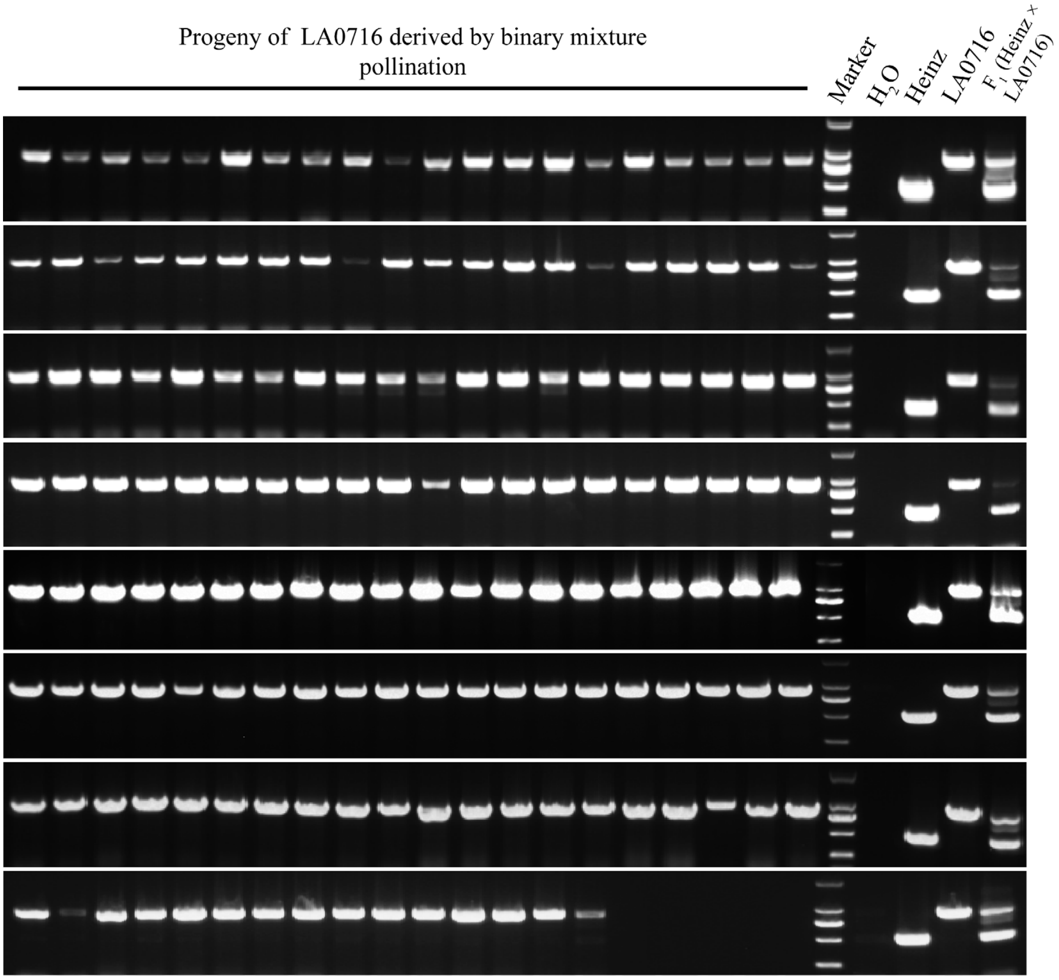


**Fig. S1** The Heinz genotype was not been detected in 155 progeny of LA0716 derived by binary mixture pollination. Genotyping was conducted using an InDel marker of *CUL1* (Li & Chetelat, 2010).

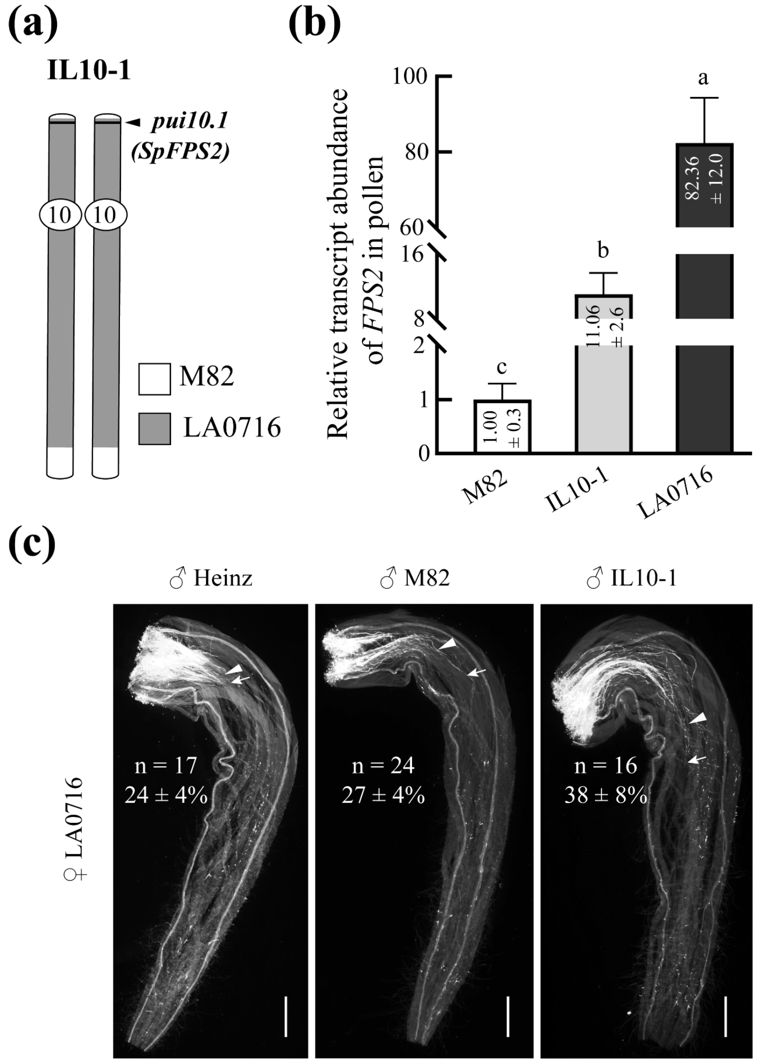


**Fig. S2** UI response remains intact in introgression line 10-1 (IL10-1) harboring pollen UI factor *SpFPS2* in LA0716 styles.

**(a)** Chromosome schematic diagram of IL10-1. The *pui10.1* identified in this study is labeled by a black bold line on Chr. 10, which contains the *pui* factor *SpFPS2* (Qin *et al.*, 2018). Introgressed fragments from LA0716 are shown in grey and chromosomal segments of M82 are shown in white. A scale representing physical length of these three chromosomes is shown. **(b)** Relative transcript abundances of *FPS2* in pollen of M82, IL10-1 and LA0716 examined by RT-qPCR. Data are presented as means ± SD (one-way ANOVA (LSD test), $P < 0.05$, n = 4). **(c)** Aniline blue staining showing pollen tube growth of Heinz, M82 and IL10-1 in LA0716 styles 72 h after pollination, respectively. White triangles and white arrows represent a majority of pollen tubes and the longest pollen tubes in pistils, respectively. Scale bar, 1 mm.

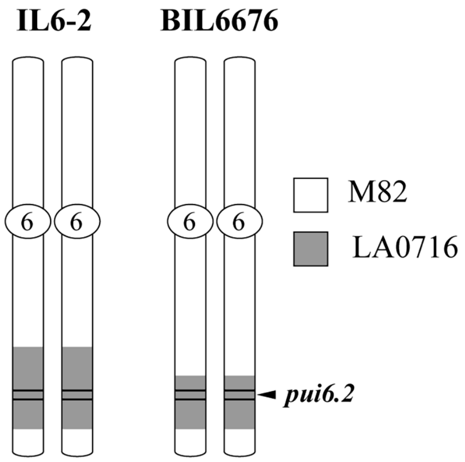


**Fig. S3** Chromosome schematic diagram of IL6-2 and BIL6676.

The *pui6.2* identified in this study is in the region labeled by two black lines on Chr. 6. Introgressed fragments from LA0716 are shown in grey and chromosomal segments of M82 are shown in white. A scale representing physical lengths of these three chromosomes are shown. IL6-2 exhibits a strong necrotic phenotype in leaves and stems, which makes it difficult to grow to reproductive stage. Thus, only BIL6676 was used in subsequent experiments.

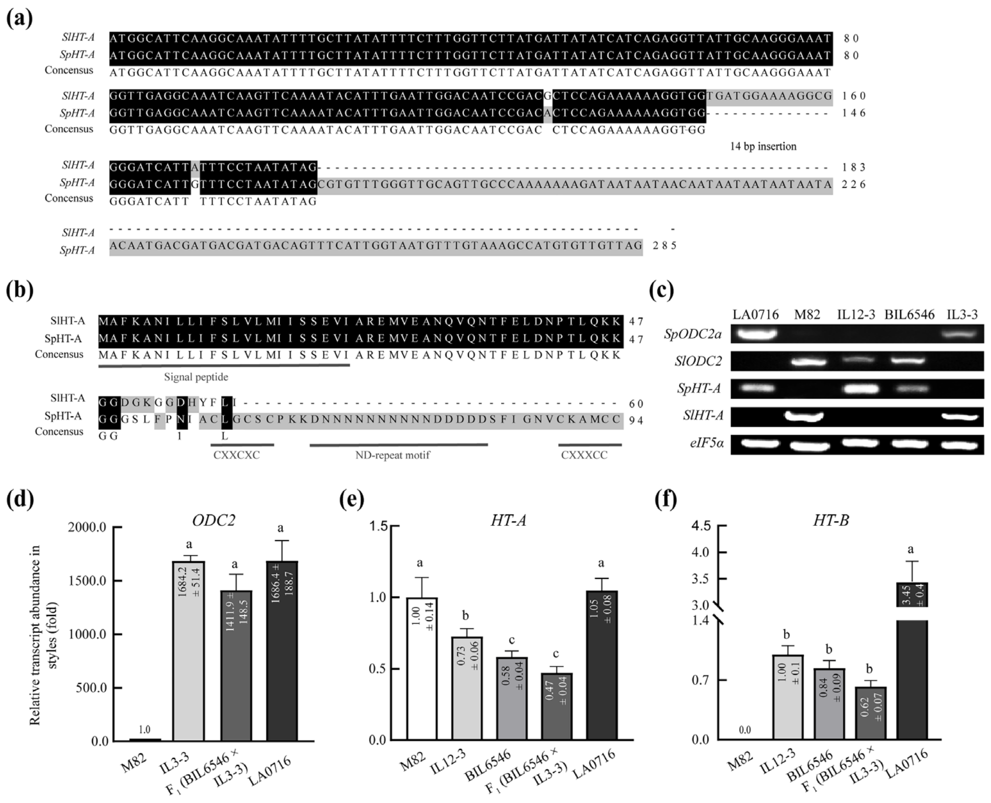


**Fig. S4** Functional comparison of stylar UI factors between M82 and corresponding ILs/BILs.

Sequence alignment of nucleotide **(a)** and amino acid **(b)** of *HT-A* genes from *S. pennellii* and *S. lycopersicum*. Compared to *SpHT-A*, 14-base pairs are inserted in the 2$^{nd}$ exon of *SlHT-A*, which results in the loss of C-terminus. C-terminus of SpHT-A contains two cysteine-rich motifs (CXXCXC and CXXXCC) and a ND-repeat motif as shown in **(b)**. **(c)** Genotypes of *SpODC2*, *SlODC2*, *SpHT-A* and *SlHT-A* in corresponding materials were analyzed with specific primers (**Table S3**). Relative transcript abundances of *ODC2* **(d)**, *HT-A* **(e)** and *HT-B* **(f)** in corresponding materials examined by RT-qPCR. Data are presented as means ± SD (one-way ANOVA (LSD test), $P < 0.05$, n = 3).

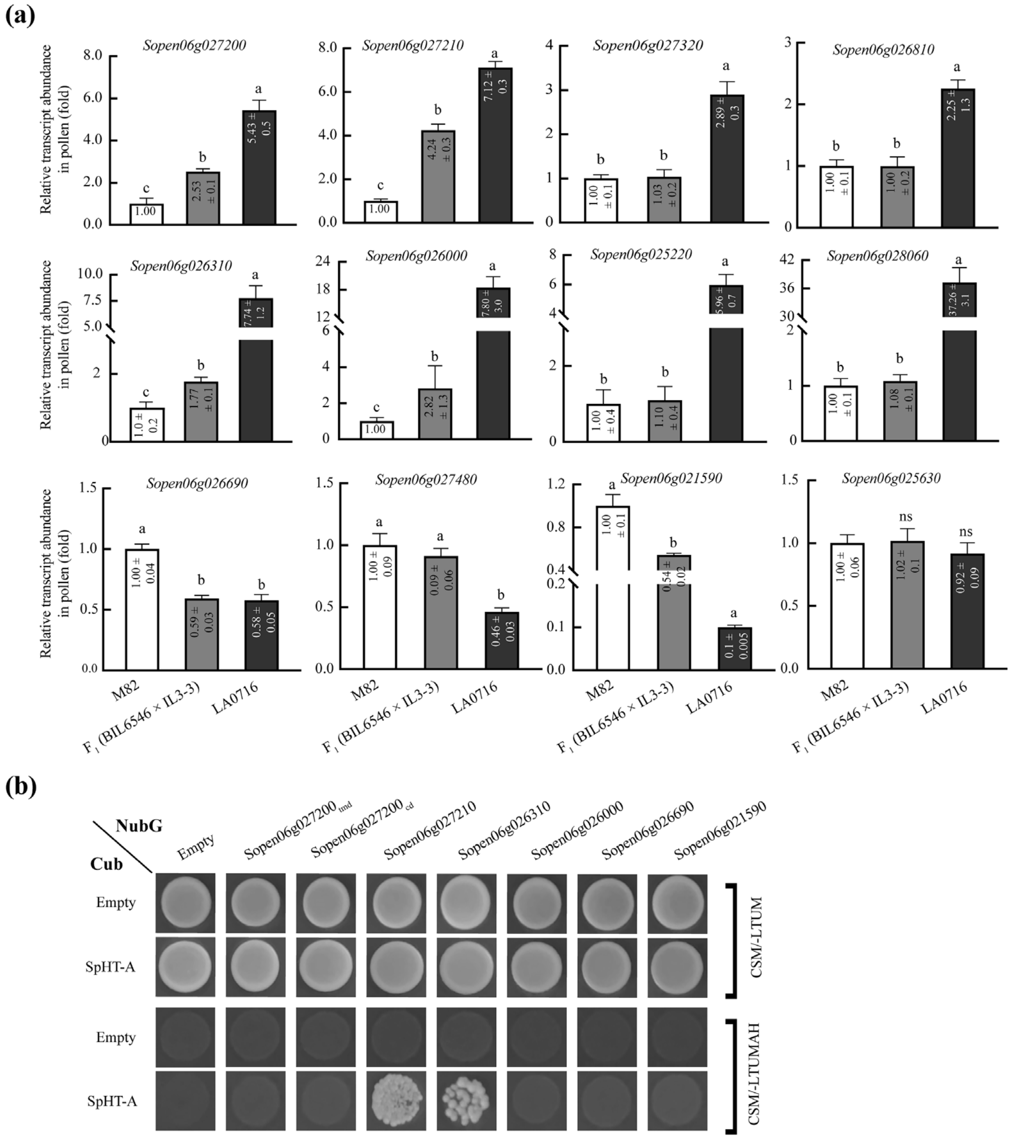


**Fig. S5** Among candidate genes linked to *pui6.2*, significant differences in *Sopen06g027210* and *Sopen06g026310* transcript abundances between $F_1$ (BIL6546 × IL3-3) and M82 pollen, and Y2H interactions with SpHT-A were found.

**(a)** Relative transcript abundance of candidate gene in pollen of M82, LA0716 and $F_1$ (BIL6546 × IL3-3) examined by RT-qPCR. Data are presented as means ± SD (one-way ANOVA (LSD test), $P < 0.05$, n = 3). Genes with significant differences in transcript abundance in pollen between $F_1$ (BIL6546 × IL3-3) and M82 were selected for further validation. **(b)** mbSUS-based Y2H of interactions between 6 candidate proteins and SpHT-A, respectively. Since neither *Escherichia coli* nor yeast with plasmid containing full-length of *Sopen06g027200* could grow on medium, its transmembrane domain (Sopen06g027200$_{tmd}$) and cytoplasmic domain (Sopen06g027200$_{cd}$) were separately cloned into the prey vector to verify their interactions with SpHT-A.

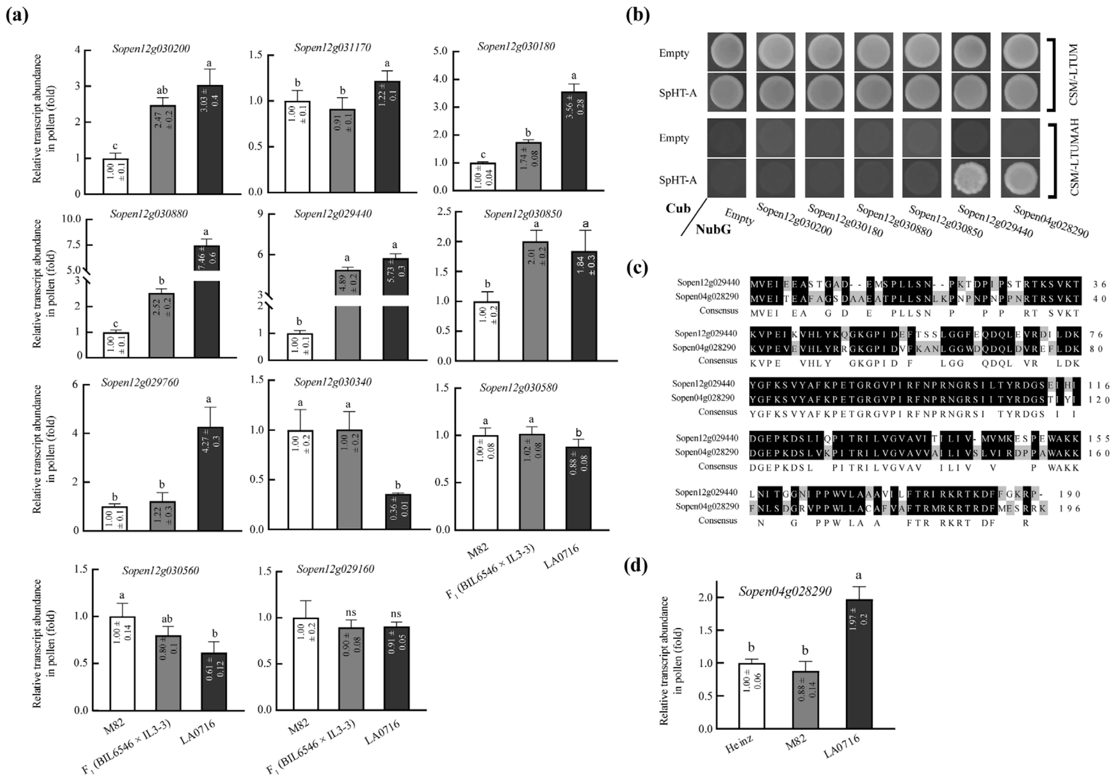


**Fig. S6** Among candidate genes linked to *pui12.1*, significant difference in *Sopen12g029440* transcript abundance was found in pollen between $F_1$ (BIL6546 × IL3-3) and M82 pollen, and both its protein product and homologous protein encoded by *Sopen04g028290* directly interacted with SpHT-A in yeast.

**(a)** Relative transcript abundances of candidate genes in pollen of M82, LA0716 and $F_1$ (BIL6546 × IL3-3) examined by RT-qPCR. Data are presented as means ± SD (one-way ANOVA (LSD test), $P < 0.05$, n = 3). Genes with significant difference in transcript abundance in pollen between $F_1$ (BIL6546 × IL3-3) and M82 were selected for further validation. **(b)** mbSUS-based Y2H analysis of interactions between 6 potential candidate proteins with SpHT-A. **(c)** Amino acid sequence alignment of two homologous WD repeat proteins encoded by *Sopen12g029440* and *Sopen04g028290.* **(d)** Relative transcript abundance of *Sopen04g028290* in pollen of Heinz, M82 and LA0716 examined by RT-qPCR.

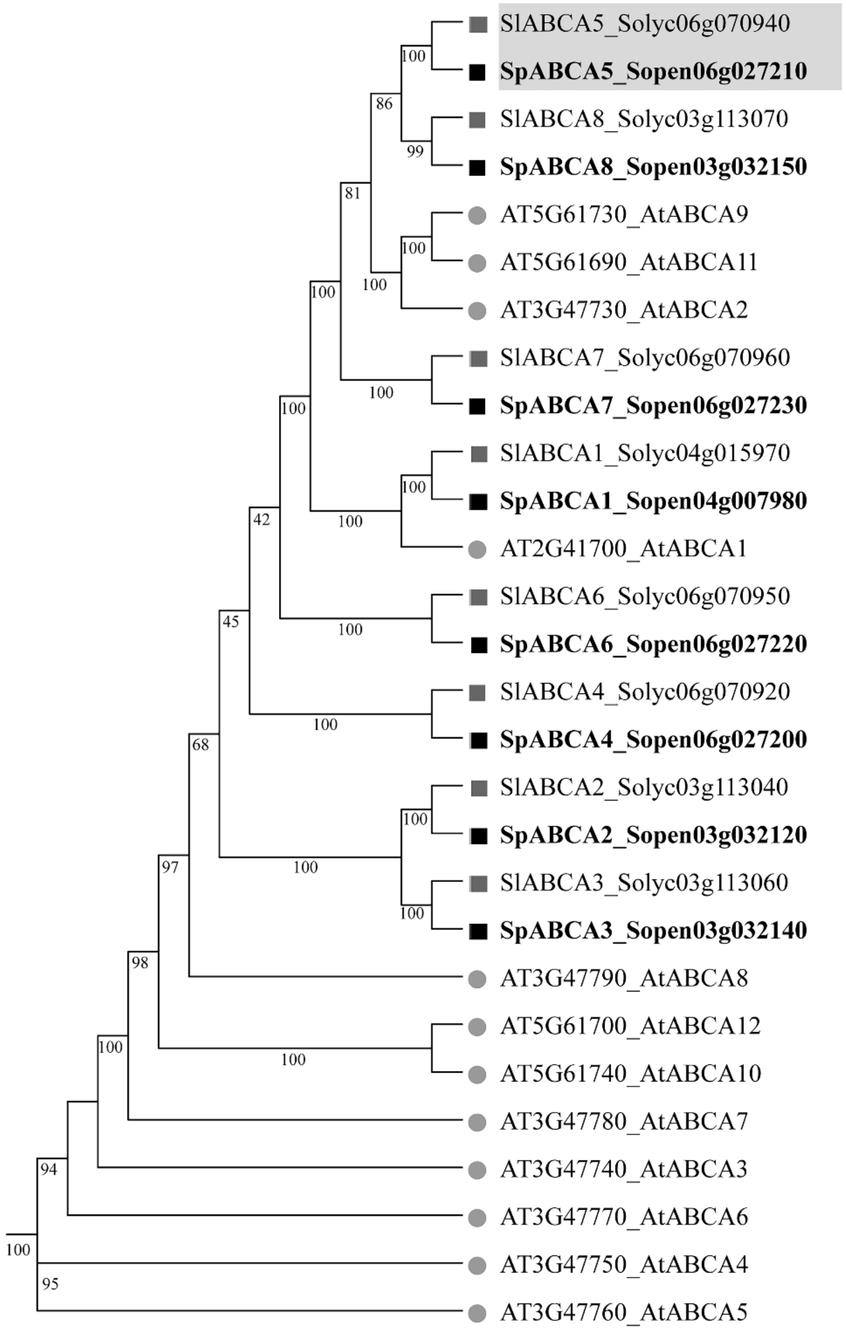


**Fig. S7** Phylogenetic tree of ABC-A subfamily proteins of *S. pennellii*, *S. lycopersicum* and *Arabidopsis thaliana*. 8, 8 and 12 ABC-A proteins were identified in genomes of *S. pennellii*, *S. lycopersicum* and *Arabidopsis thaliana*, respectively. Numbers on branches indicate the bootstrap percentage values calculated from 1000 replicates. The candidate gene *Sopen06g027210* and its orthologous gene *Solyc06g070640* (*SlABCA5*) were labeled by a rectangular gray shade.
